# Benchmarking zero-shot single-cell foundation model embeddings for cellular dynamics reconstruction

**DOI:** 10.64898/2026.03.10.710748

**Authors:** Xueya Zhou, Zihan Wang, Yue Ling, Qinxue Tian, Zhenyi Zhang, Yongge Li, Luonan Chen, Peijie Zhou

## Abstract

Reconstructing cellular trajectories from time-resolved single-cell transcriptomics is fundamental to understanding processes from embryonic development to cancer progression. While single-cell foundation models (scFMs) promise universal biological representations through large-scale pretraining, their capacity to capture the non-linear dynamics governing cell-fate decisions remains uncharacterized. Here we systematically benchmark multiple scFMs across challenging biomedical scenarios involving branching lineages and continuous state transitions. By coupling zero-shot scFM embeddings with dynamic optimal transport, we evaluated their performance against a traditional highly variable gene (HVG) baseline in backtracking progenitor states, interpolating transition intermediates, and extrapolating future fates. We find that zero-shot scFM embeddings underperform the HVG baseline across diverse biological systems, particularly in recovering the distributional complexity of unobserved cells. Mechanistic analysis reveals that current scFM architectures tend to over-compress subtle temporal signals, causing an artificial “linearization” of branched biological structures that may obscure critical divergence points in disease progression. Our findings suggest that while scFMs provide unified cell-state views, the HVG baseline remains more robust for trajectory inference, identifying a fundamental “temporal-compression” bottleneck that must be addressed to develop next-generation, dynamics-aware foundation models.

## Introduction

Understanding how cells change state over time in response to development, differentiation, or external perturbations is a central problem in biology^1^. High-throughput single-cell profiling has enabled the characterization of molecular states across vast cellular populations, providing detailed snapshots of transcriptional heterogeneity^1,3–6^. However, these assays are destructive, the same cell cannot be observed as it changes over time or in response to perturbations. Consequently, time-series single-cell experiments yield a sequence of unaligned snapshots rather than direct longitudinal measurements. Reconstructing continuous cellular dynamics from such snapshots therefore requires computational strategies that (i) embed sparse, noisy, high-dimensional single-cell transcriptomes into informative low-dimensional representations, and (ii) estimate transportation maps across time to approximate population-level flows.^7,8^

Traditionally, cell embeddings for trajectory analysis are obtained by selecting highly variable genes and applying dimensional-reduction methods such as principal component analysis, after which lineage structure and cellular dynamics are inferred in the resulting low-dimensional space^9^. In parallel, single-cell foundation models like Geneformer^2^, scGPT^10^, Genecompass^11^, are pretrained on large collections of scRNA-seq profiles to learn general-purpose representations of cells and genes, which are then used in a zero-shot manner for downstream analyses. These models have been reported to transfer to diverse tasks, including cell clustering, annotation, batch correction, and gene-level applications such as regulatory network inference^2,10–18^. Because they are exposed to a wide range of tissues, conditions, and species during pretraining, their embeddings are expected to capture shared structure in gene-expression space, to be robust to technical variation, and to support data-efficient analysis in new settings^15^. If such properties extend to temporal processes, single-cell foundation models might provide more informative representations for trajectory inference than conventional HVG-based embeddings, by better preserving subtle transitional states and stabilizing estimates of population-level flows across time. Nonetheless, existing benchmark studies have primarily focused on conventional, largely static single-cell analysis tasks, such as cell clustering, annotation, and batch correction, and have shown that, in zero-shot settings, single-cell foundation models often provide limited or inconsistent gains over simple HVG-based baselines^19–21^. In contrast, the performance of these models on explicitly dynamical tasks has been much less explored. Recent benchmarks on perturbation prediction suggest that foundation models do not necessarily outperform traditional approaches in this setting^22–25^, while their utility for reconstructing cellular dynamics and trajectories from snapshot time-series data has not yet been systematically evaluated. It therefore remains unclear whether embeddings produced by such foundation models actually confer an advantage over an HVG-based baseline for reconstructing cellular dynamics from snapshot time-series data.

Cell dynamic processes can be modeled computationally using trajectory inference methods, which order cells along a trajectory based on similarities in their expression patterns^6^. Many widely used approaches like graph-based methods (e.g. Monocle^26^) reconstruct branched manifolds from neighborhood graphs, and RNA velocity based methods (e.g. scVelo^27–30^) infer local transcriptional dynamics for individual cells based on unspliced and spliced mRNA. However, these approaches do not explicitly model the coupling between successive sampling times. In contrast, optimal transport(OT) based approaches (e.g. Waddington-OT^31^) are explicitly formulated for time-series designs, in which distinct cell populations are sampled at successive time points^8,32–34^. Given embeddings of cells at two or more times, optimal transport estimates couplings that describe how the population at an earlier time is probabilistically mapped into the population at a later time, thereby providing a principled framework for reconstructing population-level flows from destructive snapshot measurements^8,31,35–38^. In single-cell biology, OT has been used to align distributions across developmental stages or treatment time courses and to recover trajectories and fate biases in systems such as reprogramming, differentiation, cancer development and response to perturbation^38^. Extensions based on unbalanced OT^39–42^ allow total mass to vary between time points and have been introduced to better accommodate proliferation, death, and population expansion or contraction dynamically^37^. However, single-cell transcriptomes are high-dimensional and sparse, and directly fitting OT in raw expression space is statistically unstable and computationally demanding^8^. Consequently, OT-based analyses are typically performed on low-dimensional embeddings obtained by selecting HVG and applying dimensionality reduction methods, and the inferred dynamics depend critically on this embedding choice^43^. This dependence motivates a systematic benchmark of whether embeddings learned by single-cell foundation models provide advantages over simple HVG-based baselines for OT-based reconstruction of cellular dynamics from time-series data.

Although promising zero-shot transfer has been reported for general single-cell analysis tasks, the suitability of scFM embeddings for trajectory inference and dynamical reconstruction remains unresolved. To address this gap, a systematic benchmark was conducted to test whether zero-shot embeddings from five published scFMs improve trajectory inference relative to a baseline constructed from highly variable genes projected with principal component analysis (HVG-PCA) across a diverse range of complex biological systems, including hematopoietic lineage branching, embryoid body development, the epithelial-to-mesenchymal transition (EMT) and the directed in vitro differentiation of stem cells into pancreatic βcells. We evaluated three canonical scenarios that address fundamental inquiries in developmental and disease biology: backtracking seeks precursor or progenitor states that precede observed samples^44^; interpolation aims to recover transient intermediates that connect adjacent sampled time points^45^; and extrapolation anticipates downstream states beyond the last observation under continued progression of the process^46^. And performance was quantified with complementary metrics: (i) distribution recovery; (ii) pseudotime correlation; (iii) local velocity coherence^47–49^. This benchmarking design isolates embedding choice from downstream modeling, enabling a direct assessment of whether foundation model embeddings preserve the intricate non-linearities of cellular dynamics or introduce structural biases that could misinform the design of engineered biological systems.

## Results

### Overview of benchmark framework

To assess whether zero-shot embeddings from single-cell foundation models (scFMs) provide advantages for trajectory inference, a benchmark was designed that isolates representation quality from downstream modeling. Given the sparsity and dimensionality of single-cell expression data, performing trajectory inference directly in the original gene expression space is both computationally challenging and sensitive to noise. Consequently, in practice, cellular dynamics are typically inferred after projecting cells into a lower-dimensional representation, which makes the choice of embedding a central determinant of downstream trajectory reconstruction^7^. As illustrated in **Fig. 1a**, time-stamped snapshot expression profiles were first mapped by each scFM, as well as an HVG-PCA baseline, into low dimensional cell embeddings. Trajectory inference was then performed directly in this embedding space to reconstruct cellular dynamics, thereby separating the effect of representation learning from that of downstream dynamical inference.

**Fig. 1.**
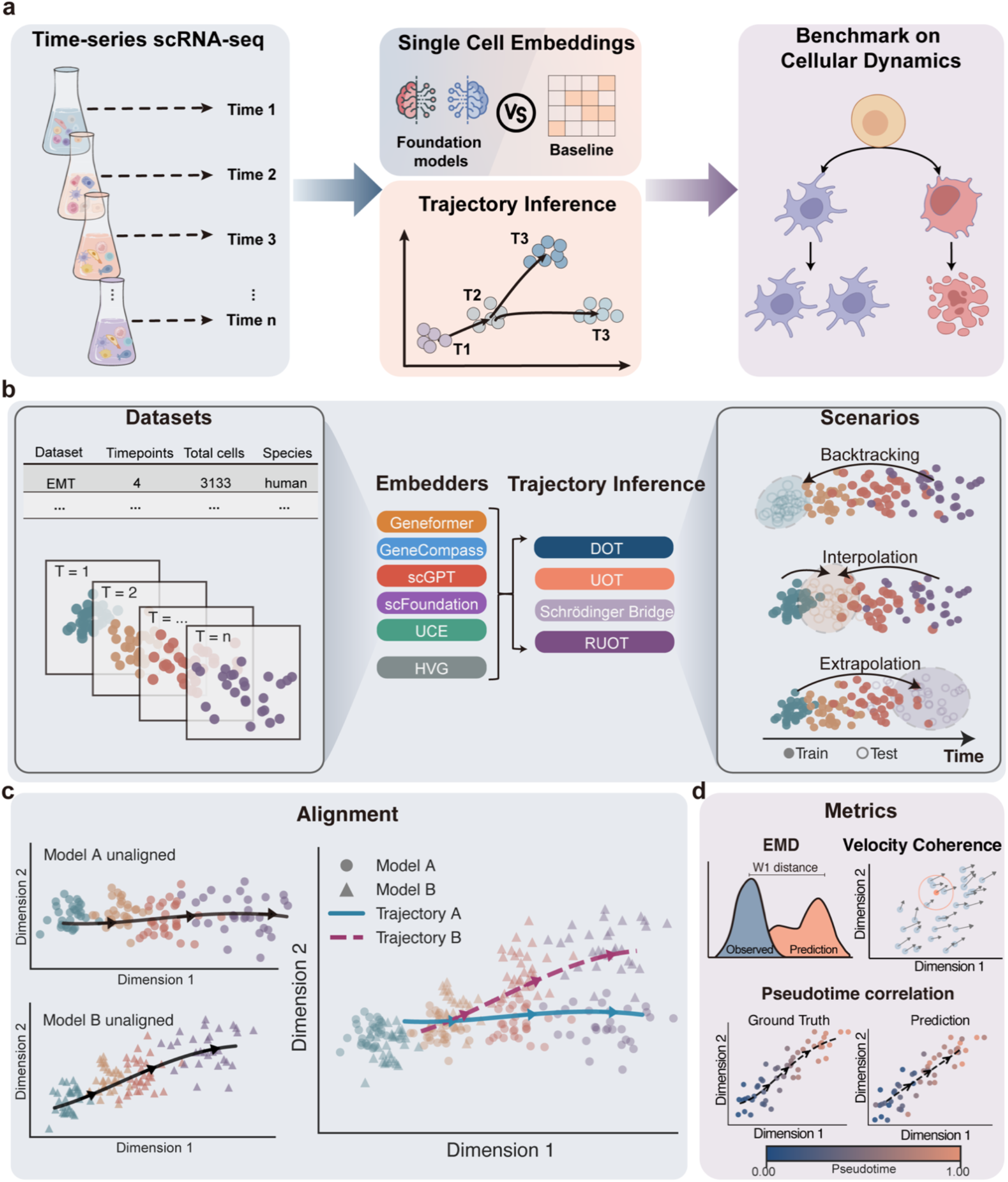
Benchmark workflow and evaluation pipeline for reconstruction of cellular dynamics in foundation model embedding space. **a**, Overview of the workflow. Single-cell transcriptomic snapshot data are embedded by large pretrained models to obtain cell representations. Trajectory inference is subsequently performed in the embedding space to reconstruct cellular dynamics. **b**, Detailed benchmark design. Snapshot data are embedded by five foundation models and one baseline method (i.e. PCA embedding with highly variable genes) to generate time-resolved cell embeddings. Dynamic optimal transport (DOT), Unbalanced Dynamical Optimal Transport (UOT), Dynamical Schrödinger Bridge, and Regularized Unbalanced Optimal Transport (RUOT), four dynamic inference methods are then applied. Data are partitioned into training and test sets based on sampling time points to simulate three dynamical tasks: backtracking, interpolation, and extrapolation. **c**, For fair comparison, embeddings and inferred trajectories from different models are aligned into a unified latent space, in which all performance metrics are computed. **d**, Illustration of the three evaluation metrics used in this study. Wasserstein-1 distance (W1) quantifies the distributional divergence between predicted and observed cell state distributions at held-out time points. Velocity coherence measures local directional consistency of inferred dynamics by computing the agreement of velocity vectors among neighbouring cells: for a representative cell (highlighted in orange), velocity vectors of its *K* nearest neighbours (circled) are compared using cosine similarity. Pseudotime correlation is defined as the Spearman rank correlation between the inferred pseudotime ordering and the reference pseudotime.

Our benchmark spans multiple published time-series snapshot datasets (Methods), covering a range of experimental systems and scales, from approximately 3,000 to 49,000 cells (**Fig. 1b**, a complete list of datasets is provided in **Supplementary Table 1**), encompassing fundamental processes such as differentiation, development, pathological transitions and cellular reprogramming. Each dataset is processed by six embedding pipelines: five foundation model embedders (Geneformer, Genecompass, scGPT, UCE, and scFoundation) and an HVG-based baseline. On top of these representations, we apply four trajectory inference methods that are closely related through optimal transport formulations and their entropy-regularized variants, including Dynamical Optimal Transport (DOT), Unbalanced Dynamical Optimal Transport (UOT), Dynamical Schrödinger Bridge^50,51^, and Regularized Unbalanced Optimal Transport (RUOT)^37,52–54^, yielding a comprehensive set of “embedding × inference” combinations for systematic comparison.

To evaluate temporal generalization, we partition each dataset by sampling time points and construct three complementary tasks (**Fig. 1b**, right). In backtracking, models are fit on later time points and used to reconstruct initial states. In interpolation, intermediate time points are held out and predicted from other time points. In extrapolation, models are fit on early time points and evaluated on later, unseen time points (Methods). This design mirrors practical scenarios with incomplete temporal coverage and tests whether embeddings and inference methods can recover cellular dynamics beyond the observed time window.

Because different embedders and trajectory inference methods generate representations in distinct coordinate systems, we align both the cell embeddings and inferred trajectories into a shared latent space prior to evaluation (**Fig. 1c**)^55^. This alignment normalizes for coordinate differences, ensuring that performance comparisons reflect recovered dynamical structure rather than representational arbitrariness. Within this shared space, performance is quantified using three complementary metrics that capture the distributional, directional, and ordinal aspects of cellular dynamics (**Fig. 1d**, Methods). Specifically, we employ the Wasserstein-1 distance (Earth Mover’s Distance, EMD) to quantify discrepancies between predicted and observed cell state distributions at held-out time points (distributional); a velocity coherence score to assess the local consistency of inferred velocity vectors (directional); and a pseudotime correlation, defined as the Spearman correlation between inferred pseudotime and the reference chronological order (ordinal). Together, these metrics provide an integrated assessment of each pipeline’s ability to reconstruct cell state distributions, flow directions, and temporal progression.

### HVG outperform zero-shot foundation model embeddings across the majority of tasks and metrics

To evaluate whether zero-shot embeddings from single-cell foundation models provide advantages for optimal transport (OT) based trajectory reconstruction beyond a simple HVG–PCA baseline, we applied the benchmark framework described above to six embedders in combination with four trajectory inference methods across multiple time-series single-cell datasets. In the aligned evaluation space, HVG–PCA embeddings yield the strongest distributional recovery across tasks and datasets. As summarized by the Wasserstein-1 (EMD) distance (lower is better; **Fig. 2a**), HVG– PCA attains the lowest discrepancies between predicted and observed held-out cell-state distributions in backtracking, interpolation, and extrapolation settings. For each dataset–task combination, points represent the mean performance across the four inference methods, with whiskers indicating the corresponding range. Among the foundation models, Geneformer and scGPT are the most competitive, but both remain inferior to the HVG baseline, whereas scFoundation, based on direct expression value encoding (Methods), shows the weakest performance. This suggests that traditional HVG representations better preserve the multi-modal heterogeneity inherent in complex transitions, whereas foundation model embeddings may suffer from representation collapse, particularly in the challenging regimes of backtracking and extrapolation where the model must infer progenitor states or future disease states beyond the observed window.

**Fig. 2.**
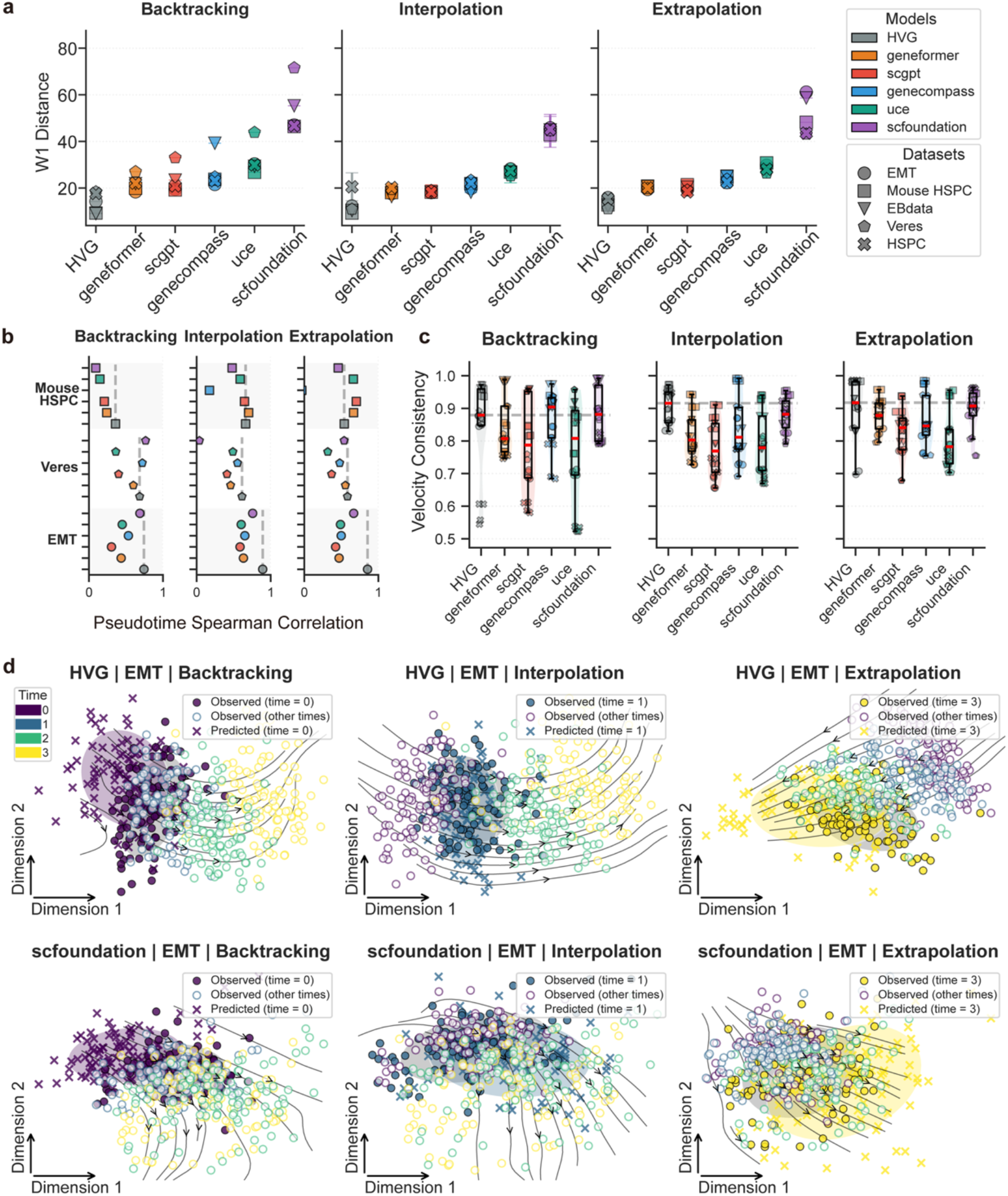
Zero-shot embeddings underperform the highly variable gene (HVG) baseline across three dynamical reconstruction tasks. **a**, Distribution of Wasserstein-1 (W_1_) distances between predicted and ground-truth cell distributions, evaluated on all datasets for backtracking, interpolation, and extrapolation. Lower W_1_ values indicate more accurate recovery of the true distributions. **b**, Spearman correlation between inferred pseudotime and reference pseudotime provided by the original dataset publications, computed across all tasks and datasets. Higher correlation coefficients reflect stronger agreement with temporal trajectories. **c**, Local velocity coherence across datasets and tasks. Scores range from 0 to 1, with higher values denoting greater consistency among velocity vectors within local cell neighborhoods. **d**, Trajectory reconstructions on the EMT dataset for the best-performing embedding (HVG) and the worst-performing embedding (scFoundation). Six subplots are organized into two rows—top row: HVG; bottom row: scFoundation—and three columns corresponding, from left to right, to backtracking, interpolation, and extrapolation. In each subplot, true cells are shown as circles, predicted cells as “×” markers, and inferred velocity vectors are overlaid as streamlines.

Temporal ordering accuracy, quantified by the Spearman correlation with reported pseudotime, exhibits a similar overall pattern. Because pseudotime lacks an absolute ground truth, correlations are computed against the reference pseudotime provided in the original data publications after alignment in the shared latent space (Methods). As shown in **Fig. 2b**, the HVG–PCA baseline achieves the highest pseudotime correlation on the EMT dataset, reaching a Spearman’s ρ of 0.892 under the interpolation setting. In datasets with more complex temporal and branching structures (EBdata and HSPC), inspection of the DPT^56^ pseudotime annotations reveals weaker concordance with experimental sampling time (**Supplementary Fig. 1**). Specifically, pseudotime values are not uniformly distributed across sampling time points and do not exhibit a clear monotonic increase with progression. In these settings, pseudotime correlation is therefore treated as a secondary metric and interpreted alongside distributional and velocity-based measures.

Directional consistency further supports this overall trend. Velocity coherence quantifies the local smoothness of inferred dynamics by measuring agreement among velocity vectors within cell neighborhoods (Methods). As shown in **Fig. 2c**, velocity coherence (higher is better) is reported across tasks for each dataset and trajectory inference method; points correspond to individual inference methods, while the red line indicates the median value across datasets and embeddings. Overall, HVG–PCA achieves the highest velocity coherence across the majority of datasets and task regimes, indicating that OT-based trajectory inference tends to recover smoother and more self-consistent dynamical flows in HVG–PCA space. Notably, in the more challenging backtracking and extrapolation settings, certain foundation model embeddings, particularly GeneCompass and scFoundation exhibit comparable performance to HVG–PCA and, in some datasets, achieve the highest velocity coherence. These results suggest that while HVG–PCA provides the most robust substrate for directional consistency overall, specific foundation models can be competitive under particular dynamical regimes.

To provide qualitative intuition for these quantitative differences, we visualize inferred trajectories on the EMT dataset in **Fig. 2d**. In a two-dimensional PCA projection of the embedding space, predictions at held-out time points substantially overlap the observed cell populations for HVG–PCA across all task settings, effectively reconstructing unobserved cell state in EMT process. In contrast, trajectories inferred in the scFoundation embedding space show limited overlap and fail to reconstruct the withheld distributions to a comparable extent. Taken together, these results indicate that, under zero-shot settings, HVG–PCA embeddings provide a more reliable substrate for trajectory inference than current single-cell foundation model embeddings, with the largest performance gaps observed in distributional accuracy and local directional coherence, and with backtracking and extrapolation representing the most challenging regimes.

### Sensitivity analyses confirm robustness to alignment strategy, reference space, and latent dimensionality

Because our benchmark integrates heterogeneous embeddings and reconstructs cellular dynamics in a reduced latent space, it is important to assess whether the observed performance differences depend on specific alignment choices or modeling hyperparameters. We therefore conducted a series of sensitivity analyses to evaluate the robustness of our conclusions with respect to embedding alignment, reference space selection, and latent dimensionality.

We first examined the role of embedding alignment, which is required to place representations produced by different models which are often differing in scale, orientation, and dimensionality into a common coordinate system prior to evaluation. Using the EMT extrapolation task as an illustrative example (**Fig. 3a**), alignment of HVG, Geneformer, and scFoundation embeddings to a consensus latent space substantially reduces inter-embedding differences in scale and orientation, enabling direct comparison of inferred trajectories and predicted cell state distributions. Aligned embeddings for all datasets are visualized in **Supplementary Fig. 2–6**. In contrast, analyses performed in the original, unaligned embedding spaces are confounded by pronounced scale mismatches and rotations, which impair comparability and can bias downstream metrics.

**Fig. 3.**
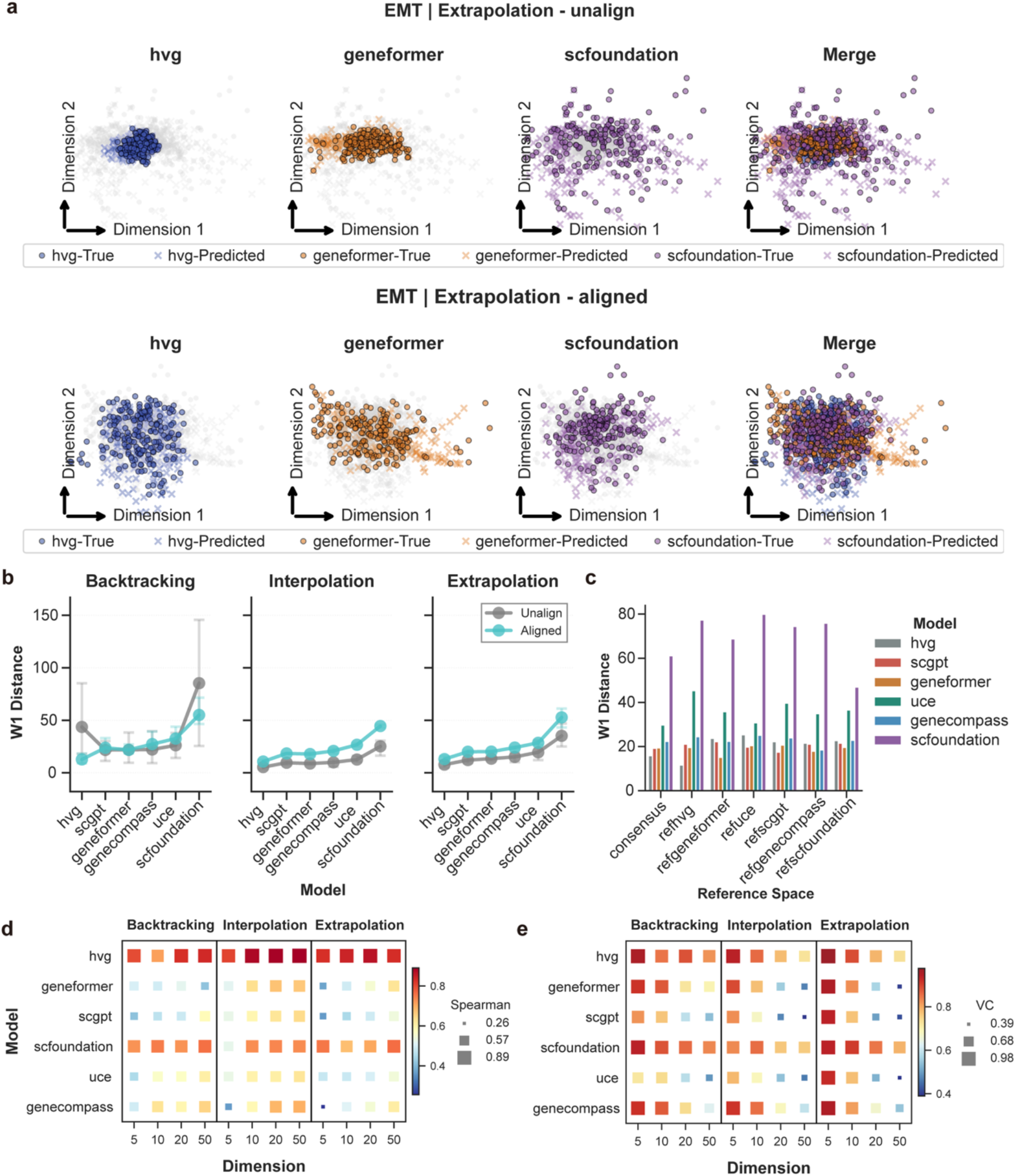
Sensitivity analyses with respect to alignment, reference space, and embedding dimensionality. **a**, Visualization of the EMT dataset illustrating evaluations with and without alignment. For HVG, Geneformer, scFoundation, and an integrated “merge” representation (combining model-specific embeddings), observed cell states and predicted states are displayed as scatter plots. The aligned condition is shown in the top row and the unaligned condition in the bottom row; columns correspond to the four representations. Observed and predicted states are distinguished by marker shape. **b**, Wasserstein-1 distance evaluated across the three dynamical tasks under alternative alignment settings. “Unalign” denotes evaluation in each model’s native embedding space without alignment. “Aligned” denotes orthogonal Procrustes alignment to a reference embedding, corresponding to an orthonormal rotation and reflection that minimizes squared distances between embeddings. **c**, Effect of the choice of reference space on Wasserstein-1 distance. Results are reported after mapping embeddings into different reference spaces; “consensus” denotes a shared latent space obtained by aggregating embeddings aligned across all models. **d**, Heatmaps showing the effect of embedding dimensionality on pseudotime correlation across the three dynamical tasks. The horizontal axis indicates the number of retained principal components. Color encodes the Spearman correlation between inferred pseudotime and the reference pseudotime, with blue indicating lower and red indicating higher correlation (values approaching 1 indicate stronger agreement). **e**, Heatmaps showing the effect of embedding dimensionality on local velocity coherence across the three tasks. The horizontal axis indicates the number of retained principal components. Color encodes coherence, with blue indicating lower and red indicating higher values (values approaching 1 indicate stronger local directional consistency of velocity vectors).

Having established the necessity of alignment, we next examined whether the benchmark conclusions depend on the specific alignment strategy or choice of reference space. Comparing aligned and unaligned evaluations reveals that the absence of alignment introduces modest shifts in Wasserstein-1 distances and can alter the relative ranking of embedders in some cases (**Fig. 3b**), underscoring the importance of embedding alignment for fair comparison. Nevertheless, when alignment is applied, varying the alignment procedure yields stable relative rankings of embedders by distributional accuracy, as measured by the Wasserstein-1 distance. To further assess potential reference-space bias, we repeated the interpolation task while alternately treating each embedding as the reference space for alignment (**Fig. 3c**). Although absolute W1 values shift modestly, which often favoring the embedding chosen as the reference, the overall performance ordering across embedders remains largely unchanged. Consistent trends are observed for pseudotime correlation and velocity coherence under different alignment and reference space choices (**Supplementary Fig. 7**), indicating that the robustness of our conclusions is not specific to a single evaluation metric. Based on these sensitivity analyses, all main results are reported using Generalized Procrustes Analysis (GPA) alignment to a consensus latent space (Methods)^57^.

We additionally assessed sensitivity to a key optimal transport hyperparameter: the dimensionality of the latent space used to learn cellular dynamics. Across latent dimensions of 5, 10, 20, and 50, pseudotime correlations preserve the overall ranking of embedders (**Fig. 3d**), although the optimal dimensionality varies across embeddings. For example, in the Genecompass embedding space, a higher-dimensional latent representation substantially outperforms lower-dimensional alternatives, indicating that appropriate dimensionality selection can materially improve performance within a given embedding.

Velocity coherence exhibits a similar pattern of robustness at the cross-embedding level (**Fig. 3e**), with relative performance largely preserved across latent dimensionalities. Within individual embeddings, lower dimensional latent spaces generally yield higher local velocity coherence, consistent with the recovery of smoother and more locally self-consistent flow fields on a more compact manifold. Sensitivity analyses across all datasets are provided in **Supplementary Fig. 8**.

Taken together, these sensitivity analyses demonstrate that our central findings i.e. the superior performance of HVG-based embeddings over zero-shot foundation model embeddings across most tasks and evaluation metrics, are robust to the choice of alignment strategy, reference space, and latent dimensionality. This robustness supports the use of the benchmark as a general and reliable framework for evaluating embedding representations in dynamic single cell settings.

### Zero-shot foundation model embeddings attenuate temporal and branching structure relevant for trajectory inference

To investigate why zero-shot foundation model embeddings exhibit systematic limitations on dynamics tasks, we examined how they represent temporal structure. We defined a time variance ratio (TVR), the fraction of total variance attributable to time (Methods), to quantify the extent to which time-scale differences are preserved. As illustrated by a linear discriminant analysis on the EMT dataset (**Fig. 4a**)^58^, foundation model embeddings exhibit markedly reduced TVR relative to the HVG baseline, indicating diminished temporal separability. Pairwise energy distances between time points, visualized as a heatmap, further support this observation: time points exhibit increased proximity and reduced discriminative resolution in foundation model spaces than in the HVG representation. Among the foundation models, scFoundation retains comparatively greater temporal separation, although still below HVG and without translating into consistent improvements across downstream dynamics metrics.

**Fig. 4.**
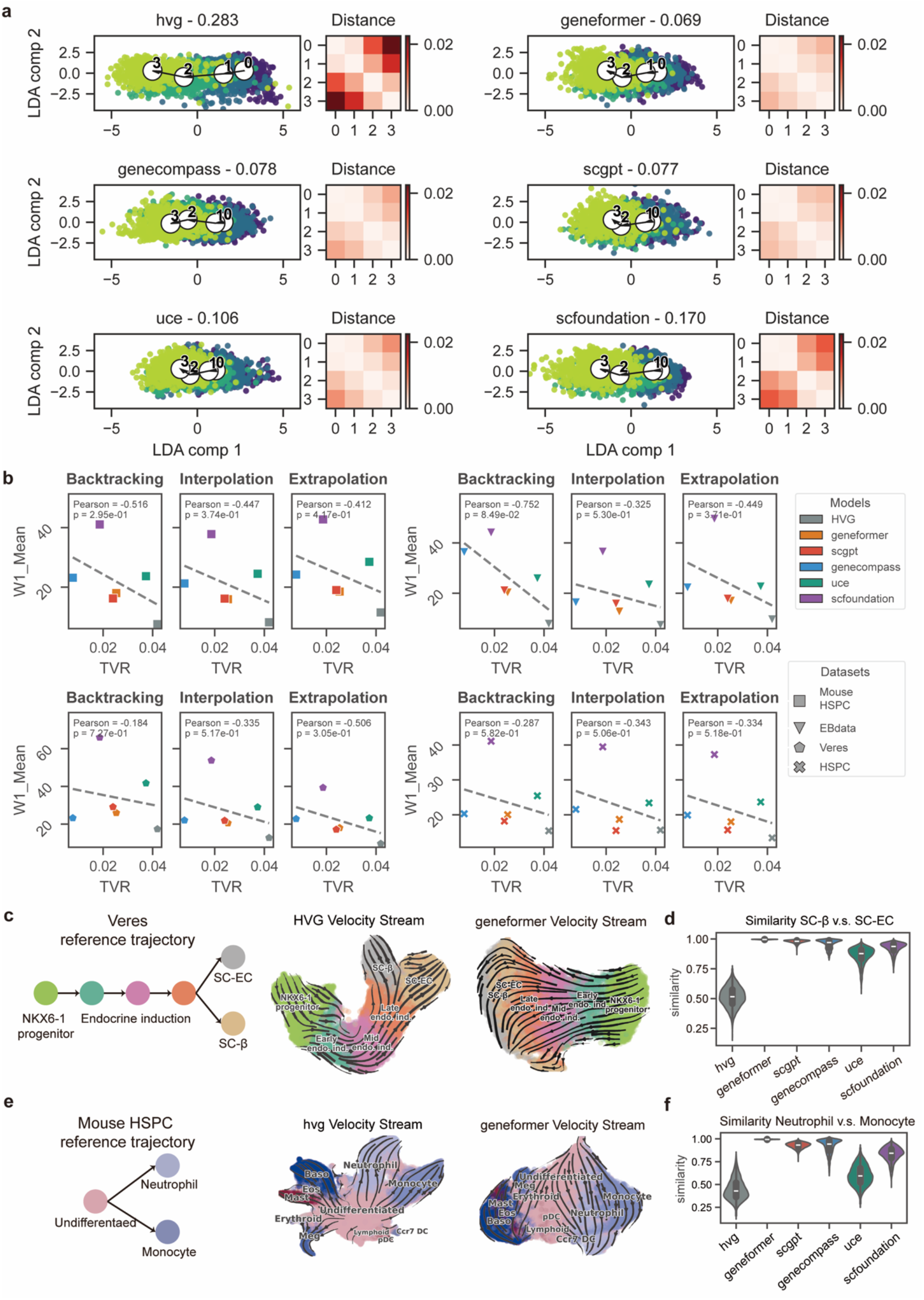
Batch correction like temporal compression and impaired branch resolution in large-model embeddings and their association with diminished performance in dynamical reconstruction. **a**, EMT dataset: for each embedding model, an LDA visualization colored by sampling time and a heatmap of pairwise energy distances between time points are shown; panel titles report the Temporal Variance Ratio (TVR, higher indicates greater temporal separation). Reduced TVR together with attenuated between–timepoint distances indicates temporal compression. **b**, Association between temporal separation and reconstruction error across four datasets and three tasks. Each scatter plot relates TVR to the Wasserstein-1 distance (W1); points are colored by embedding model. A negative association is observed, indicating that lower temporal separation is linked to higher distributional error. **c**, Human pancreatic differentiation (Veres dataset). Left: reference trajectory depicting differentiation from NKX6-1^+^ progenitors toward SC-β and SC-EC fates. Middle: HVG embedding visualized with UMAP and extrapolation velocity streamlines inferred by RUOT (regularized unbalanced optimal transport). Right: Geneformer embedding visualized with UMAP and RUOT extrapolation streamlines. In the HVG embedding, SC-β and SC-EC branches remain well separated and extrapolated flows follow the reference branches; in the Geneformer embedding, these fates are partly merged and extrapolated flows become less branch-aligned. **d**, Pairwise similarity between SC-β cells and SC-EC cells across embeddings from different single-cell foundation models (scFMs), quantified by cosine similarity. **e**, Mouse HSPC. From left to right: reference trajectory; HVG embedding with UMAP visualization and RUOT extrapolation velocity streamlines; Geneformer embedding with UMAP visualization and RUOT extrapolation velocity streamlines. **f**, Pairwise similarity between Neutrophil and Monocyte populations across scFM embeddings.

We next examined the relationship between temporal variance preservation and dynamics performance. Across four datasets and three dynamical scenarios (**Fig. 4b**), higher TVR is generally associated with lower Wasserstein-1 distances, indicating improved recovery of held-out cell-state distributions when time-scale differences are preserved. However, this negative association is not universal. For example, in the EMT dataset (**Supplementary Fig. 9**), Geneformer, scGPT, and GeneCompass display very low TVR, with time points nearly co-localized in the embedding space, yet achieve low W1 distances. In this regime, predicting held-out distributions becomes comparatively trivial because observed and target states are already close in the embedding. Similar analyses relating TVR to pseudotime correlation and local velocity coherence in multiple datasets (**Supplementary Fig. 10, 11**) further support that compression of time-structured variation is generally accompanied by degraded dynamical reconstruction.

Motivated by this pattern, we asked whether foundation model embeddings might be attenuating temporal signal in a manner analogous to batch-effect correction. In the EMT dataset, explicitly applying Harmony^59^ batch correction similarly reduces between time variance and equalizes W1 across embeddings, but at the cost of lower pseudotime correlation and velocity coherence (**Supplementary Fig. 12**). This controlled perturbation supports the view that removing time-structured variation, whether by design or implicitly in foundation model embeddings, tends to impair dynamical reconstruction.

We then asked whether a similar “compression” occurs along branching, fate related axes. In the Veres human pancreatic differentiation dataset, HVG embeddings preserve two clear branches from NKX6-1^+^ progenitors toward SC-β and SC-EC fates (**Fig. 4c**, left), and extrapolated RUOT trajectories follow these reference branches in a fate-specific manner (**Fig. 4c**, middle). By contrast, in the Geneformer embedding, SC-β and SC-EC populations are less clearly separated, and extrapolated flows are less tightly aligned with the two terminal branches (**Fig. 4c**, right). Consistently, cosine similarity analyses show increased similarity between SC-β and SC-EC states in multiple foundation model embedding spaces (**Fig. 4d**). An embedding incorporating protein sequence information during pretraining (UCE) exhibits comparatively improved separation between these two endocrine populations, but does not yield consistently better dynamical performance. A similar pattern is observed in mouse hematopoiesis (**Fig. 4e, f**), in this bifurcating system, large-model embeddings increase the similarity between Neutrophil and Monocyte populations relative to HVG, and RUOT-based velocity streamlines become less branch-aligned in the large-model space than in the HVG space, indicating that branch-specific biological variation is systematically compressed in zero-shot foundation model embeddings.

Overall, temporal and branching analyses together indicate that zero-shot foundation model embeddings systematically compress both time-scale differences and branch-specific biological variation. Such compression of biologically meaningful structure, likely arising from an over-correction of variation treated as nuisance (e.g. batch-like effects), offers a plausible explanation for the consistently weaker performance of downstream dynamical methods in large-model embedding spaces relative to HVG-based embeddings.

### Global benchmarking reveals consistent advantages of HVG-based embeddings across methods and datasets

To synthesize the benchmark results across all evaluated settings, we next summarized dynamical performance for all combinations of embedding spaces, trajectory inference methods, tasks, and datasets using an integrated heatmap representation (**Fig. 5**). This global view enables simultaneous assessment of consistency and variability across the full benchmarking landscape.

**Fig. 5.**
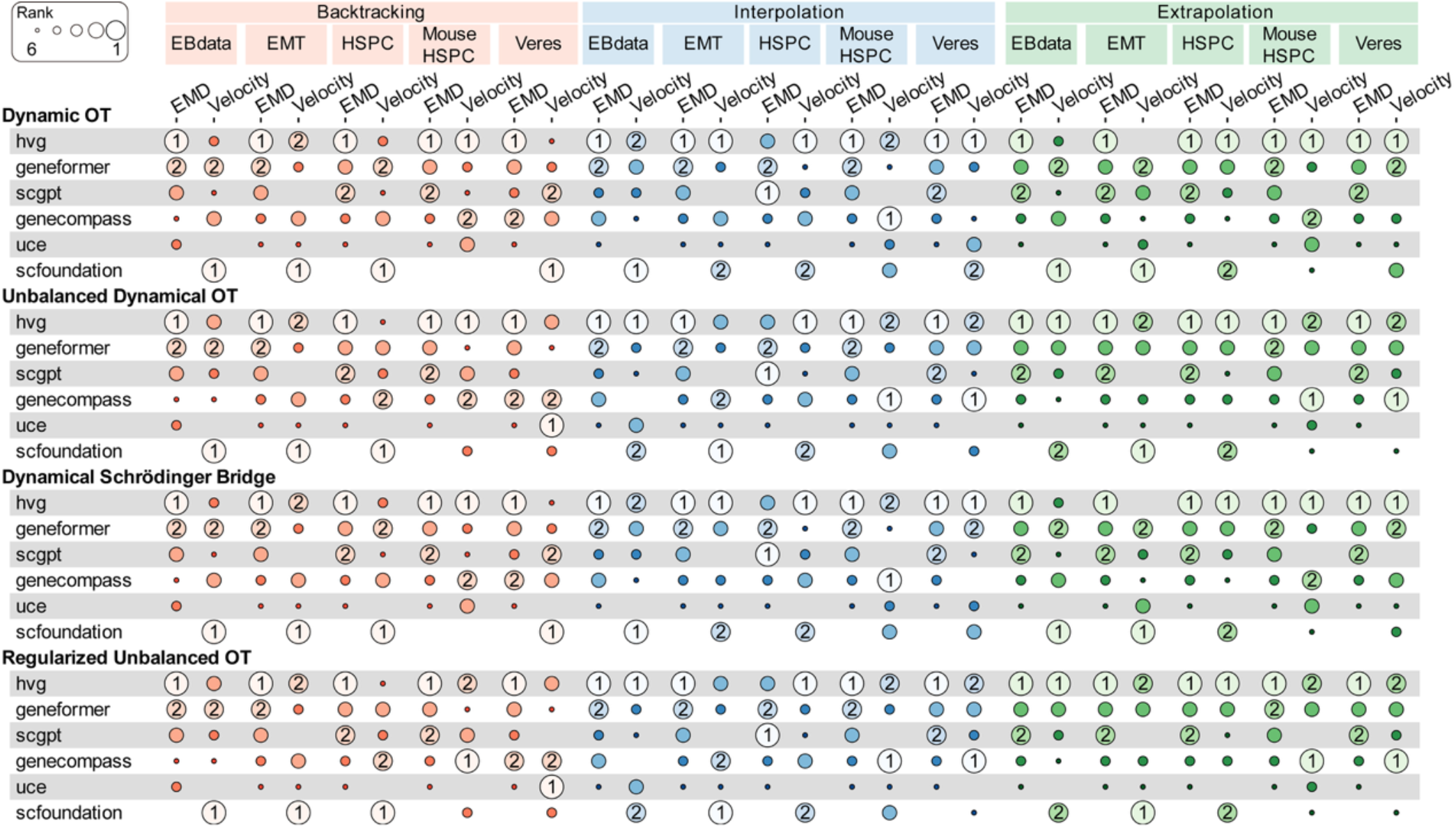
Global rank-based summary of embedding performance across datasets and tasks. Heatmap showing the relative ranking of embedding methods across all datasets, three dynamical tasks (backcasting, interpolation, and extrapolation), and optimal-transport–based trajectory inference settings. For each dataset– task–inference combination, embeddings are ranked based on distributional accuracy (EMD) and local velocity coherence, with higher ranks indicating better relative performance. For interpolation tasks involving multiple intermediate test time points, ranks are aggregated across these settings. The top two performing embeddings for each dataset–task combination are highlighted. Heatmaps of the corresponding raw metric values are provided in the Supplementary Information (Supplementary Note 3).

Across the majority of settings, the HVG-based baseline exhibits superior performance relative to zero-shot single-cell foundation model embeddings, most prominently in distributional accuracy as measured by the EMD. This advantage is observed consistently across datasets, task regimes, and optimal-transport–based inference methods, indicating that the superiority of HVG-based representations is not driven by a particular modeling choice or experimental system. Among foundation models, Geneformer generally achieves stronger distributional recovery, whereas scFoundation tends to yield higher local velocity coherence in some settings; however, no single foundation model embedding consistently dominates across all metrics or tasks.

In **Fig. 5**, we summarize global benchmark performance using rank-based heatmaps rather than raw metric values. For each dataset, task, and trajectory inference method, embeddings are ranked according to distributional accuracy (Wasserstein-1 distance) and local directional consistency (velocity coherence), and these ranks are then aggregated to provide a comparative overview across settings. This rank-based representation emphasizes relative performance trends and reduces the influence of dataset specific scale differences across metrics. Heatmaps of the corresponding raw metric values for all settings are provided in the **Supplementary Note3, Fig13, Fig14**.

Taken together, this global analysis confirms that the performance advantages of HVG-based embeddings over zero-shot foundation model embeddings are robust across inference methods, datasets, and task settings. At the same time, the heterogeneity revealed by the heatmap highlights that current foundation models exhibit metric- and task-specific strengths, suggesting that future improvements will require embedding strategies that better preserve temporal and biological-specific structure rather than relying on zero-shot representations alone.

## Discussion

In this study, a systematic benchmark was conducted to compare single-cell foundation models (scFMs) with a highly variable gene (HVG) baseline for cellular dynamics reconstruction across three tasks: backtracking, interpolation, and extrapolation. All methods were evaluated in a shared aligned embedding space using complementary metrics that capture different aspects of dynamical reconstruction: (i) distributional recovery, quantified by the Wasserstein-1 distance (Earth Mover’s Distance), (ii) global agreement with reference pseudotime, quantified by Spearman correlation, and (iii) local velocity coherence, which measures neighborhood-level consistency of inferred velocity vectors.

Based on their training paradigm and architecture, scFMs pretrained with transformer models on large single-cell corpora would be expected to capture gene–gene dependencies and higher order structure through self-attention, thereby encoding information relevant to cellular dynamics and potentially outperforming a simple HVG baseline. However, contrary to this expectation, we found that the HVG-based embedding consistently outperformed scFM embeddings in most settings and across metrics. This pattern was robust to the choice of alignment procedure and latent dimensionality. Among scFMs, Geneformer and scGPT achieved comparatively stronger performance on the Wasserstein-1 metric, whereas scFoundation yielded higher local velocity coherence. Pseudotime correlation appeared to be more sensitive to data topology: in datasets with bifurcating trajectories, several scFM embeddings tended to linearize branches, thereby obscuring biologically meaningful branching structure while artificially inflating pseudotime correlations. Beyond intrinsic data geometry, our analyses also indicated that zero-shot scFM embeddings can over correct batch effects, reducing temporal separability and degrading performance on dynamical tasks. Taken together, these results show that, in their current zero-shot form, scFM embeddings are not superior to a straightforward HVG-based baseline for reconstructing cellular dynamics, and may in fact suppress temporal variation by treating it as batch-like noise.

A possible explanation for these observations lies in the inductive biases introduced by current self-supervised training objectives and model architectures. Most scFMs are trained with reconstruction or masked token prediction losses that encourage invariance to technical perturbations and emphasize stable co-expression structure. Such objectives are well aligned with tasks that require robust cell identity representations, but they inherently prioritize features that are frequent and persistent across cells^60^. By contrast, transient transcriptional programs that govern short lived state transitions are less consistently sampled and therefore contribute less to the self-supervised signal. As a result, the learned embeddings are dominated by static, identity related variation (general signals), whereas temporally restricted dynamics (specific signals) are under-represented. This interpretation is consistent with recent benchmarking studies reporting that zero-shot embeddings from scFMs often underperform HVG-based baselines on tasks that depend on fine-grained or context specific variation, including perturbation responses prediction^25,60^. Together, these observations suggest that current scFMs preferentially capture general, context-agnostic structure at the expense of process-specific temporal signals.

Architectural and encoding choices likely reinforce this bias. Rank- or bin-based preprocessing of gene expression compresses dynamic range and down-weights gradual expression changes, while pooling gene token embeddings into a single cell level representation further compresses local structure on the expression manifold^18^. These design choices stabilize training and enhance invariance to nuisance variation, but they also smooth out fine-scale curvature and reduce separability between time points and branches. In developmental and perturbation trajectories, where relevant signals are often concentrated in a subset of genes and confined to specific temporal windows, such smoothing can collapse biologically distinct intermediate states into a more homogeneous representation^22,24,61^. This offers a geometric explanation for the reduced temporal variance and branch blurring observed in zero-shot scFM embeddings.

Taken together, our findings imply that general purpose scFMs, as currently trained, may be intrinsically better suited to static tasks such as cell type classification and batch integration than to reconstructing dynamical processes. This functional divergence is rooted in the model’s architecture and pretraining objectives, which we illustrate conceptually in **Fig. 6**. Current scFMs map diverse biological contexts into a relatively uniform embedding space with a collapsed geometry. By prioritizing identity-related stability, these models treat most temporal variation as noise to be removed, thereby obscuring short-lived transitional programs and directly degrading downstream performance on dynamics. Closing this gap will likely require making dynamical structure an explicit target of representation learning rather than a purely downstream objective. In particular, models will need to preserve temporal differences between states and maintain the separation of distinct developmental or perturbation trajectories. Overall, developing scFMs that explicitly balance robustness to technical variation with retention of biologically meaningful temporal structure represents a key avenue for future work.

**Fig. 6.**
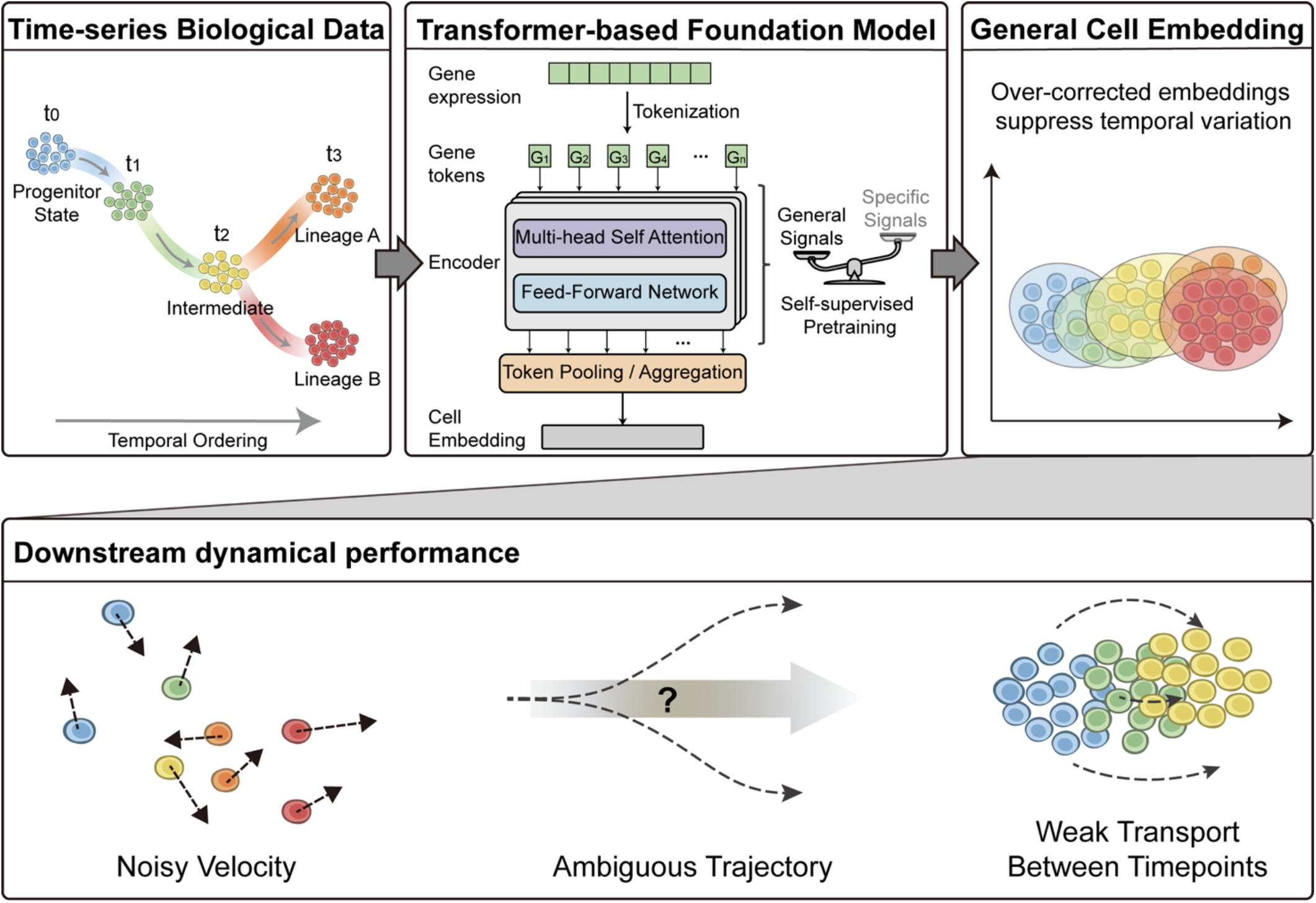
Conceptual understanding of temporal information loss in scFM embeddings: Inductive biases in current single-cell foundation models (scFMs) prioritize stable cell identity over transient dynamical signals. Input (Left): Time-resolved biological data with structured trajectories and distinct branching lineages (t0 – t3). Mechanism (Center): Self-supervised pretraining introduces an inductive bias (represented by the balance) that favors persistent “General Signals” (e.g., cell type) while down weighting transient, process specific “Specific Signals”. Embedding (Right): The resulting latent space exhibits collapsed geometry, where temporal variation is suppressed and distinct cellular states overlap. Impact (Bottom): Loss of fine-grained temporal information degrades downstream dynamical tasks, manifested as Noisy Velocity vectors, Ambiguous Trajectories, and Weak Transport between time points.

## Methods

### Benchmark framework overview

The benchmark framework comprises three core steps: (i) scRNA-seq data are reduced to a low-dimensional cell space using different embedding methods; (ii) trajectory-inference algorithms are applied in this space to model cellular dynamics; and (iii) predicted cell states and trajectories are aligned to a consensus reference space, where three evaluation metrics are computed.

### Foundation models and baseline

For all single-cell foundation models (scFMs), we first obtain cell embeddings from the model encoders. In this study, we use the [CLS] token embedding from the encoder as the representation of each cell, construct a time-series of cell embeddings with the original hidden dimension, and then apply PCA to reduce them to a low-dimensional space suitable for trajectory inference.

#### Geneformer

Geneformer V2 is a foundational transformer model pretrained on ∼104 million non-cancer human single-cell transcriptomes. It encodes each cell using a rank-based representation in which genes are ordered by relative expression and scaled using population-level statistics, emphasizing cell-state-informative genes while reducing the influence of ubiquitous housekeeping genes. For each cell, the top 4,096 ranked genes are selected as the input sequence. The encoded sequences pass through multiple Transformer encoder layers and are trained using a masked-gene prediction objective, enabling fully self-supervised learning on unlabeled data. In this study, we use the Geneformer-12L-4096i-104M model, which is pretrained on 104 million human single cells and takes 4,096 ranked genes as input, encoded by a 12-layer Transformer encoder.

#### Genecompass

Genecompass is a knowledge-informed cross-species foundation model self-supervised pre-trained on an extensive dataset scCompass-126M, which comprises over 120 million high-quality human and mouse single-cell transcriptomes. For each cell, the model takes the top 2048 ranked genes after normalization and ranking of gene expression values, and incorporates four types of prior biological knowledge (including GRN, promoter sequence, gene family and co-expressions) encoded within a unified embedding space. It employs a 12-layer Transformer architecture and adopts a masked language modeling objective which randomly masks 15% of gene inputs in each cell.

#### UCE

UCE is a cross-species single-cell foundation model, which employs a 33-layer Transformer encoder with 650 million parameters. It is pre-trained on over 36 million cells spanning eight species in a completely self-supervised way without any data annotations. Gene embeddings are initialized using the pretrained protein language model ESM-2^62^, incorporating protein biological knowledge, genes are sampled based on expression probability and sorted by chromosomal order; the pretraining task uses masked binary prediction, training the model to determine whether a gene is expressed in a cell. This enables UCE to establish a unified biological latent space capable of representing any cell, regardless of tissue or species. In this study, we use the 33-layer UCE model to obtain cell embeddings.

#### scFoundation

scFoundation is a large-scale pretrained model with 100M parameters, which models 19,264 genes and is pre-trained on over 50 million scRNA-seq data. It adopts an asymmetric Transformer-like encoder-decoder architecture xTrimoGene, whose embedding module converts continuous gene expression values into high-dimensional learnable vectors rather than using discretized values, and employs an asymmetric encoder-decoder structure to handle the high sparsity of single-cell data, enabling efficient learning of all gene relationships.

#### scGPT

scGPT is a single-cell foundation model for multiple single-cell omics data analysis applying large-scale pretrained Transformer. The model takes 1,200 HVGs as input and introduces a cell-dependent value binning technique to discretely encode gene expression levels. It employs a special attention masking mechanism to adapt the Transformer to non-sequential omics data while incorporating condition tokens to encode metadata, enabling it to simultaneously learn context-aware embeddings for both cells and genes. The pre-trained scGPT can be optimized to achieve superior performance across diverse downstream applications through fine-tuning. Here we apply the version whole-human model pretrained on 33 million normal human cells.

#### HVG baseline

As a traditional baseline, raw transcript counts are library-size normalized to 1 × 10^5^ and log1p-transformed using Scanpy v1.11.3 [ref. ^63^]. The top 2,000 highly variable genes are then selected, and PCA is applied to obtain a low-dimensional space for trajectory inference.

### Trajectory inference methods

To build a comprehensive framework for benchmarking scFM embeddings from cellular dynamics, we apply four OT-based methods to reconstruct cell-state transitions from time-series scRNA-seq datasets. The core idea of optimal transport relies on finding a mapping that minimizes a specific cost functional when transporting a probability mass from an initial distribution to a terminal state. In the context of temporally resolved snapshots data, this problem can be interpreted as an evolution of cell density *ρ*(*x, t*) governed by a continuity equation and energy minimization principle. This section provides a brief review of four typical OT-based trajectory inference methods.

The four methods considered are: Dynamical Optimal Transport (DOT), Unbalanced Dynamical Optimal Transport (UOT), Dynamical Schrödinger Bridge, and Regularized Unbalanced Optimal Transport (RUOT). In practice, all four approaches were implemented using the Python package DeepROUT^53^, with method-specific behavior controlled primarily through two parameters: *sigma*, which modulates the strength of entropy or Tikhonov regularization, and *use_mass*, which specifies whether to enforce mass conservation (balanced OT) or allow for unbalanced transport. Adjusting these parameters allows us to flexibly switch between standard, regularized, and unbalanced OT formulations while maintaining a unified computational framework across all embedding spaces and tasks.

### Dynamical Optimal Transport (DOT)

Dynamical optimal transport (DOT) is a landmark work which models cell development process as a deterministic transport characterized by *dX*_*t*_ = *v*(*X*_*t*_, *t*)*dt* [ref. ^64^]. Mathematically, this framework involves finding the velocity field *v*(*x, t*) and cell density *ρ*(*x, t*) that minimize the total transport cost:

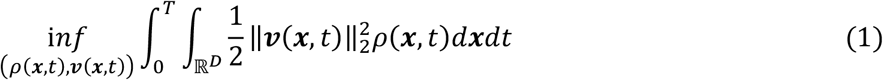

with *v*(*x, t*) and *ρ*(*x, t*) subject to the continuity equation constraint and boundary conditions:

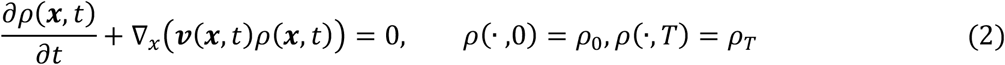

However, the strict conversation assumption in equation (2) fails to consider mass changing which is popular in biological systems due to cell proliferation and apoptosis.

### Unbalanced Dynamical Optimal Transport (UOT)

To address the limitation of mass conservation, unbalanced dynamical optimal transport (UOT) introduces a growth term *g*(*x, t*) that represents cell growth or death rate, and rewrite equation (2) as:

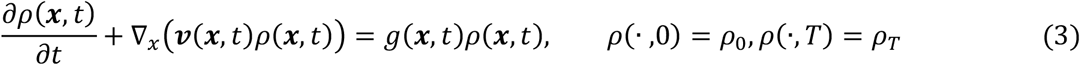

The Wasserstein–Fisher–Rao (WFR) distance is often used to quantify the overall shipping cost with respect to both kinetic and growth energy:

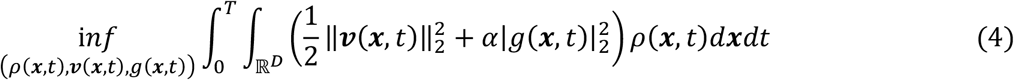

where *α* is a hyperparameter to balance the penalty of growth against transport. While UOT successfully models the unbalanced cellular dynamics, it remains a deterministic transport model that neglects the intrinsic stochasticity in gene expression and cell differentiation.

### Dynamical Schrödinger Bridge

In this case, biological processes is modelled as SDE dynamics characterized by *dX*_*t*_ = *v*(*X*_*t*_, *t*)*dt* + *σ*(*X*_*t*_, *t*)*d****W***_*t*_, where ***W***_*t*_ is a standard Brownian motion and *σ*(*x*_*t*_, *t*) denotes the random fluctuations in the system. The Schrödinger Bridge problem seeks to identify the most likely evolutionary path between an initial distribution *ρ*. and a terminal distribution *ρ*_-_, which can be formulated as solving 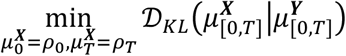 with the reference process defined as *d****Y***_*t*_ = *σ*(***Y***_*t*_, *t*)*d****W***_*t*_. This problem can be equivalently transformed into a dynamical form:

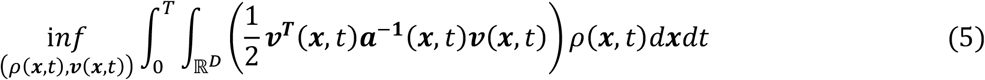

where ***a***(*x, t*) = *σ*(*x, t*)*σ*^***T***^(*x, t*) and all pairs (*ρ*(*x, t*), *v*(*x, t*)) satisfy following constraints:

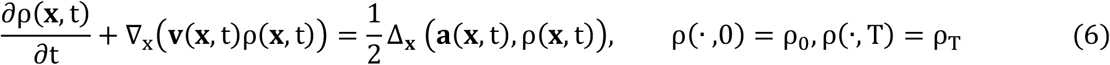

This formulation interprets trajectory inference as a stochastic optimal control problem rather than purely deterministic transport, but doesn’t incorporate unbalanced effects.

### Regularized Unbalanced Optimal Transport (RUOT)

Integrating the stochasticity in Schrödinger Bridge problem with cell proliferation effects in UOT, Regularized Unbalanced Optimal Transport (RUOT) assumes particles are governed by the SDE model *dx*_*t*_ = *v*(*x*_*t*_, *t*)*dt* + *σ*(*t*)***I****d****w***_*t*_ . RUOT seeks to minimize a generalized shipping cost accounting for velocity, diffusion, and growth penalties simultaneously:

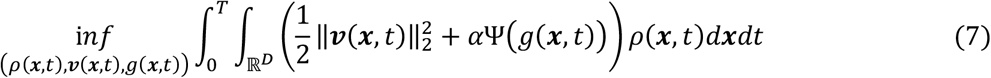

where Ψ(·) represents the growth penalty function, and all pairs (*ρ*(*x, t*), *v*(*x, t*), *g*(*x, t*)) are constrained by the unnormalized Fokker-Planck equation:

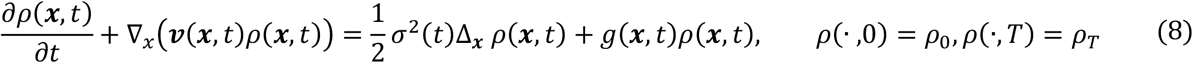

with vanishing boundary condition: 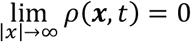

### Dynamical Reconstruction Tasks

To systematically evaluate the generalization capability of embeddings, we formulated three dynamical reconstruction scenarios: interpolation, extrapolation, and backtracking, based on the partition of sampling time points 𝒯 = {*t*_0_, *t*_1_, …, *t*_*K*_}. For each task, we hold out a specific time point *t*_test_ and train the trajectory inference models (DOT, UOT, Schrödinger Bridge, RUOT) on the remaining subset 𝒯_*train*_, subsequently inferring the cell state distribution at held-out time points.

#### Interpolation

This task evaluates the ability to recover transient intermediate states within the observed temporal window. We hold out an intermediate time point *t*_test_ = *t*_*K*_ (where 0 < *k* < *K*) and train the model using the remaining time points 𝒯_*train*_ = 𝒯 ∖ {*t*_*K*_} . The reconstruction is performed by integrating the learned dynamics starting from the initial distribution 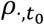 The cell state *X*_*t*_ evolves according to the forward stochastic differential equation (SDE):

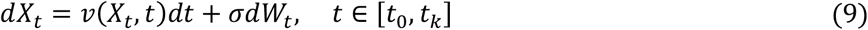

subject to the initial condition 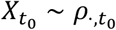 Here *W*_*t*_ denotes a standard Brownian motion. For deterministic transport, we set *σ* to 0. For unbalanced settings (UOT, RUOT), the weight of each cell is simultaneously modulated by the growth rate *g*(*x, t*):

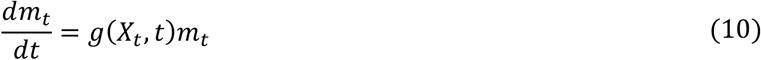

The predicted distribution 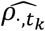 is obtained by pushing forward the population through the time interval [*t*., *t*_*K*_] and comparing it against the held-out ground truth.

#### Extrapolation

In this scenario, we assess the capacity to predict future developmental states beyond the training horizon. We hold out the last time point *t*_test_ = *tK* and train on the preceding sequence 𝒯_*train*_ = {*t*., …, *t*_*K*−1_}. Similar to the interpolation task, the inference is conducted by simulating the process starting from *t*_0_ up to the target time *t*_*K*_ . The dynamics follow the same forward SDE and growth equations defined above.

#### Backtracking

Backtracking tests the capability to reconstruct progenitor states. The model is trained on later time points 𝒯_*train*_ = {*t*_c_, …, *t*_*K*_} to predict the unobserved initial state at *t*_test_ = *t*_0_. Unlike forward tasks, this requires solving the time-reversal of the diffusion process. The reverse dynamics 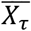, where *τ* = *T* − *t* denotes reversed time, are governed by the backward SDE:

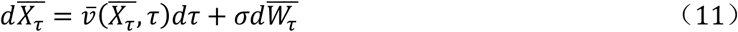

Here, the effective reverse drift 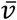 is determined by the forward velocity *v* and the score function of the density field:

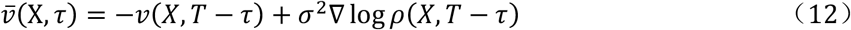

For deterministic transport (where *σ* → 0), the score term vanishes, and the reverse velocity simplifies to 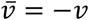. For unbalanced transport, the growth dynamics are similarly inverted. The mass evolution follows:

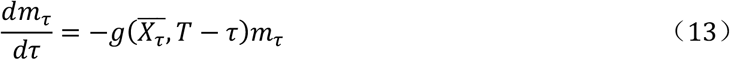

We simulate this coupled reverse process starting from the observed population at *t*_c_ to estimate the progenitor distribution 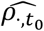

### Alignment

Alignment was used to map heterogeneous cell embeddings into a common space so that downstream metrics could be computed on comparable scales. Each aligner was fitted using cells from training time points and then applied to held-out time point. Both the reconstructed trajectories and the observed cell-expression embeddings were transformed into this common reference space before metric computation.

#### Procrustes alignment

The Procrustes problem seeks a rotation and a single global scale that best align X to Y in the least-squares sense while approximately preserving relative geometry. The optimal transformation is obtained from the singular value decomposition of the cross-covariance between X and Y, with the rotation given by the orthogonal factor and the uniform scale derived from the trace of the aligned matrices^65^. This constraint limits distortion and promotes stable, near-isometric alignment when dimensionalities are comparable.

#### Generalized Procrustes Analysis

To reduce dependence on an arbitrary reference and avoid favoring any single embedding, Generalized Procrustes Analysis (GPA) was employed to produce a consensus space. GPA iteratively (i) aligns each embedding to the current consensus using Procrustes and (ii) updates the consensus as the average of the aligned configurations; iterations continue until changes fall below a tolerance^66^. The consensus was initialized as the mean configuration across embedding spaces. The resulting per-embedding transforms, estimated on training time points, were then fixed and applied to test data. In this manner, all embeddings were compared in a reference-free, consensus-defined space.

### Evaluation Metrics

All metrics were computed in a common reference latent space to ensure comparability across methods and tasks. Under the alignment setting, both observed cells at the held-out time point and model-generated predictions were first projected into this shared space. For each dataset and task, the optimal transport (OT) trajectory model defined a time-conditioned mapping *f*(*t*) from an initial snapshot at *t*. to later states. Predictions at the held-out time *t*^∗^ were obtained by evaluating *f*(*t*^∗^) and by interpolating 100 intermediate time points between *t*. and *t*^∗^ (fixed a priori). Unless stated otherwise, the neighborhood size was K = 5 for nearest-neighbor operations.

#### Distributional recovery (Wasserstein-1 Distance)

Distributional agreement between predicted and observed cells at the held-out time point was quantified using the Wasserstein-1 distance (Earth Mover’s Distance, EMD). Let P = {*x*_*j*_}(*j* = 1 … *n*) and *Q* = {*y*_*j*_}(*j* = 1 … *m*) denote the aligned embeddings of predicted and observed cells, treated as empirical measures with uniform weights. With Euclidean ground cost *C*(*x, y*) = ‖*x* − *y*‖_2_ the distance was computed as

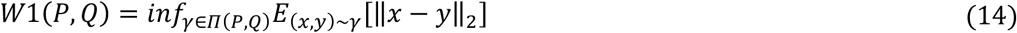

#### Pseudotime correlation

Global temporal ordering accuracy was quantified using the Spearman rank correlation between predicted and reference pseudotime values. Because pseudotime does not admit a universal ground truth, dataset-specific strategies were used to define the reference ordering. For datasets in which pseudotime annotations were provided by the original studies, these values were used directly as reference pseudotime. For datasets lacking published pseudotime labels (e.g., the Mouse HSPC dataset), reference pseudotime was estimated using diffusion pseudotime (DPT), which infers a one-dimensional progression by integrating diffusion distances along the cell-state manifold^56^. After OT fitting, 100 evenly spaced intermediate time points were interpolated between *t*. and *t*_∗_. For each observed cell, the predicted pseudotime was defined as the mean interpolation time of its K = 5 n nearest predicted neighbors among all interpolated cell states in the aligned latent space. Spearman’s ρ was then computed across observed cells between the predicted pseudotime and the reference pseudotime.

#### Local velocity coherence

Local consistency of directional change was assessed via the coherence of predicted velocities in the aligned evaluation space. After fitting the OT model, a velocity vector *v*(*X*_*t*_, *t*) was obtained for each cell, representing the predicted instantaneous direction of change at state *X*_*t*_. All generated cells at held-out time points were mapped, together with their velocity vectors, into the aligned space. For each generated cell, the K = 5 nearest generated neighbors were identified based on their positions in this space, and local velocity coherence was defined as the mean cosine similarity between the velocity of the index cell and the velocities of its neighbors, with velocity vectors normalized to unit length prior to similarity computation unless stated otherwise^49^. Higher values indicate greater agreement in the predicted directions.

### Time variance ratio

To quantify how much temporal information is encoded in each embedding space, we computed the time variance ratio (TVR) for each embedding.

For a given embedding, let *X* ∈ ℝ^n×d^ be the cell-by-dimension matrix, with cells assigned to time groups *k* = 1… *K* . We treat time as a categorical grouping variable and decompose the total variance into “between-time” and “within-time” components. We define TVR as:

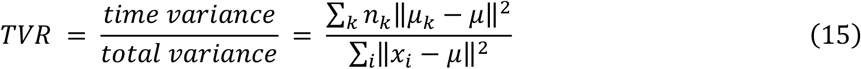

where x_*i*_ is the embedding of cell *i, μ* is the overall mean embedding, *μ*_*K*_ is the mean embedding of time group *k*, and *n*_*K*_ is the number of cells in group *k*. A higher TVR indicates that a larger fraction of the embedding variance is explained by time.

### Data collection and preprocessing

We assembled five published time-series single-cell RNA-sequencing (scRNA-seq) datasets for benchmarking (**Supplementary Table 1**), encompassing diverse biological systems, temporal resolutions, and dataset sizes (∼3,000–49,000 cells). The datasets include: EMT (GEO: GSE147405)^67^, Mouse HSPC (Weinreb et al.)^68^, Veres (GEO: GSE114412)^69^, embryoid bodies (EBdata) (Moon et al.)^70^, and HSPC (GEO: GSE226824)^71^. Raw count matrices and associated metadata were retrieved to satisfy the input requirements of single-cell foundation models (scFMs). To minimize processing-dependent artifacts in the learned representations, only minimal quality control (QC) was applied: cells with fewer than 200 detected genes and genes expressed in fewer than 3 cells were excluded. No normalization, scaling, batch correction, ambient RNA removal, or imputation was applied prior to embedding. For non-human datasets (Mouse HSPC), human–mouse one-to-one ortholog mapping was performed to enable compatibility with human-trained scFMs. Time-point annotations and lineage labels were retained as provided in the original publications.

#### EMT

This dataset captures the epithelial–mesenchymal transition (EMT) induced by transforming growth factor beta 1 (TGFB1) in the A549 cell line. Single-cell RNA sequencing was performed at five original time points (0d, 8h, 1d, 3d, and 7d). After quality control, 3,133 cells were retained for downstream analysis. To simplify trajectory modeling and ensure sufficient cell coverage at each time point, we merged the late-stage samples collected at 3 days and 7 days into a single time point, reflecting a shared late EMT state. The resulting four time points were then indexed as *t*_1_ to *t*_3_, corresponding to 0d, 8h, 1d, and the merged 3d/7d samples. This discretization was used consistently across all trajectory inference and evaluation analyses.

#### Mouse HSPC

This dataset profiles hematopoietic stem and progenitor cell (HSPC) differentiation using a combination of single-cell RNA sequencing and clonal lineage tracing, originally introduced by Weinreb *et al*. Cells were clonally barcoded using the LARRY (lineage and RNA recovery) system, enabling joint measurement of transcriptional state and lineage relationships across times. Single-cell transcriptomes were collected from heterogeneous progenitor populations at multiple time points during differentiation, both *in vitro* and following transplantation into mice. The data capture continuous hematopoietic trajectories from multipotent progenitors toward myeloid, erythroid, and lymphoid lineages, providing a well-characterized reference for evaluating trajectory inference and dynamical reconstruction methods. For compatibility with human-pretrained scFMs, one-to-one human–mouse ortholog mapping was applied prior to embedding. After filtering, a total of 49,302 cells were included in the analysis.

#### Veres

This dataset profiles in vitro differentiation of human pluripotent stem cells toward pancreatic endocrine lineages, originally reported by Veres et al. Using a high-resolution time-course single-cell RNA-sequencing design, the study captures transcriptional dynamics underlying the emergence of β-cells, α-like poly-hormonal cells, and other pancreatic and non-endocrine populations. Cells were collected across multiple stages of directed differentiation, providing temporally ordered snapshots that span early progenitor states through mature endocrine-like cell identities. For benchmarking, we adopted the original stage and time-point annotations provided in the study, and 18099 cells passed QC.

#### Embryoid bodies (EBdata)

This dataset consists of a time-series single-cell RNA-sequencing profiling of human embryonic stem cells differentiated as embryoid bodies over a 27-day period, capturing the emergence of multiple developmental lineages. Cells were collected at five sampling time points spanning early pluripotent states through diverse differentiated populations, 18,204 cells are retained after QC.

#### HSPC

This dataset characterizes the molecular and cellular dynamics of hematopoietic stem and progenitor cells (HSPCs) during inflammatory response. Single-cell RNA sequencing was performed at three time points (3, 24, and 72 hours) following treatment with the pro-inflammatory cytokine IFNα. After minimal quality control, 9983 cells were retained for analysis. Original time-point annotations and any lineage information were used as provided in the study. This dataset captures heterogeneous and dynamic transcriptional responses of HSPCs to IFNα, providing a temporal resolution suitable for benchmarking trajectory inference methods.

All datasets were subsequently processed to generate embeddings for six pipelines: five foundation model embeddings (Geneformer, GeneCompass, scGPT, UCE, and scFoundation) and a highly variable gene (HVG)-based baseline.

## Acknowledgements

We thank Professor Weinan E for the helpful discussion. This work was supported by the National Natural Science Foundation of China [Nos. 12288101 and T2321001 to P.Z., Nos. T2350003, 12131020, 42450084, 42450135, 12326614, 12426310, and T2542018 to L.C.], Zhejiang Province Vanguard Goose-Leading Initiative [No.2025C01114 to L.C.], Shenzhen Medical Research Fund [Nos. E250200620, E250200621 to L.C.], and Tianfu Jincheng Laboratory [No. TFJCPI20260001 to L.C.]. We acknowledge the support from the High-performance Computing Platform of Peking University and DP technology for computation.

## Author contributions

P.Z., L.C. and X.Z. conceived the research. X.Z. and P.Z. designed the benchmark. X.Z. and Z.W. performed benchmark and analysis. All authors discussed and interpreted the results. X.Z., Y.L. and Q.T. drafted the manuscript with the input from all authors. P.Z. and L.C. supervised the research.

