## Supplementary Information for "Benchmarking zero-shot single-cell foundation model embeddings for cellular dynamics reconstruction"

**Supplementary information**
**Supplementary Notes 1. Interpretation of pseudotime-based evaluation.**
Pseudotime correlation was included in our benchmark to assess whether the temporal ordering implied by inferred cellular dynamics is consistent with an independently defined notion of biological progression. Conceptually, a successful dynamical reconstruction should recover trajectories whose direction and progression align with established pseudo-temporal orderings derived from gene expression patterns.

However, psudotime is itself an inferred quantity and does not constitute an absolute ground truth for temporal progression. To evaluate the reliability of pseudotime annotations used in our benchmark, we therefore systematically examined their concordance with experimental sampling time across datasets (**Supplementary Fig. 1**). For datasets in which pseudotime annotations were provided by the original studies, these values were used directly as the reference ordering. For datasets lacking published pseudotime labels, reference pseudotime was estimated using diffusion pseudotime (DPT) with biologically motivated root cell selection, as described in Methods. This procedure infers a one-dimensional progression based on diffusion distances in gene expression space and is commonly used to obtain a global ordering from snapshot single-cell data.

In datasets with relatively simple or approximately linear temporal progression, pseudotime estimates exhibit a clear monotonic increase with experimental sampling time and are broadly distributed within each sampling point, consistent with coherent temporal structure. This behavior is exemplified by the EMT dataset (**Supplementary Fig. 1a**). In contrast, datasets characterized by complex branching, asynchronous differentiation, or overlapping developmental programs show substantially weaker correspondence between pseudotime and sampling time. In such cases, pseudotime values are unevenly distributed across sampling points and fail to exhibit a consistent monotonic trend, as observed for EBdata (**Supplementary Fig. 1d**).

These observations indicate that, in complex dynamical systems, pseudotime provides an ambiguous and potentially unreliable global ordering. Accordingly, in our benchmark, pseudotime correlation is treated as a secondary evaluation metric and interpreted jointly with complementary measures that more directly probe dynamical reconstruction, including distributional recovery (Wasserstein-1 distance) and local directional consistency (velocity coherence). This joint interpretation mitigates over-reliance on pseudotime in settings where a single global temporal ordering is poorly defined.

**Supplementary Note 2. Effect of explicit batch correction on dynamical reconstruction**
To further examine whether the reduced dynamical performance of zero-shot foundation model embeddings may stem from attenuation of time-structured variation, we conducted a controlled perturbation analysis using explicit batch-effect correction. This analysis was designed to test whether artificially removing between-time variance in a conventional HVG-based embedding would recapitulate the performance patterns observed in foundation model embedding spaces.

Specifically, starting from the HVG–PCA embedding of the EMT dataset, we applied Harmony-based batch correction using experimental sampling time as the batch variable. This procedure explicitly reduces variance associated with temporal progression, while preserving the underlying gene expression structure. Importantly, batch correction was applied prior to trajectory inference, and the full optimal-transport-based dynamical reconstruction pipeline was then re-fitted on the batch-corrected embeddings. All evaluations reported in this analysis focus on the interpolation setting, where intermediate time points are inferred between observed snapshots.

As shown in **Supplementary Fig. 12a**, Harmony-based correction substantially compresses between-time separation in the embedding space, yielding a representation in which cells from different sampling times are more closely co-localized. This transformation qualitatively resembles the temporal compression observed in several zero-shot foundation model embeddings. Quantitative evaluation reveals a clear metric-specific effect of batch correction on dynamical reconstruction (**Supplementary Fig. 12b**). After Harmony-based correction, Wasserstein-1 distances across embeddings are markedly flattened: predicted and observed distributions at held-out time points exhibit similar W1 values across all embeddings, reflecting that when cell states from different time points are artificially made more similar, distributional matching becomes uniformly easier. In contrast, both pseudotime correlation and local velocity coherence consistently decrease following batch correction, indicating that explicit removal of time-associated variation degrades the recovery of global temporal ordering and weakens local directional consistency of inferred dynamics. Together, these results demonstrate that while attenuating temporal differences can trivialize distributional reconstruction, it simultaneously compromises the ability to recover meaningful cellular dynamics.

Together, these results support the interpretation that suppressing time-structured variation, whether explicitly through batch-effect correction or implicitly through representation learning, can facilitate distributional matching while simultaneously impairing the recovery of biologically meaningful temporal order and dynamical flow. This controlled analysis strengthens the view that over-correction of temporal variation is one contributing factor to the weaker performance of zero-shot foundation model embeddings in dynamical trajectory inference tasks.

**Supplementary Note 3. Raw metric heatmaps underlying global rank summaries**
To complement the rank-based summaries presented in **Fig. 5**, we provide heatmaps of the underlying raw metric values across all datasets, task scenarios, embedding spaces, and trajectory inference settings (**Supplementary Figs. 13–14**). These figures report the Wasserstein-1 (W1) distance and local velocity coherence (VC) computed for each individual benchmark configuration prior to rank aggregation, where lower W1 values indicate better distributional reconstruction and higher VC values indicate stronger local directional consistency.
**Supplementary Fig. 13** summarizes the distribution of W1 distances across all datasets and three evaluation scenarios. Across nearly all settings, the HVG baseline consistently achieves the lowest W1 distances, indicating superior recovery of cell-state distributions regardless of task configuration. This trend holds across datasets and trajectory inference settings, suggesting that HVG retains strong distributional fidelity even when embeddings differ substantially in their temporal or structural properties. In contrast, local velocity coherence exhibits greater heterogeneity across datasets and task scenarios (**Supplementary Fig. 14**). While no single embedding dominates uniformly, HVG remains competitive in the majority of settings and frequently ranks among the top-performing methods. The increased variability in VC reflects the sensitivity of velocity-based metrics to local geometric structure and dataset-specific dynamical complexity.

**Supplementary Table 1 | Datasets used in benchmark.**

| Dataset | N cells | N time points | species | description | reference |
|---|---|---|---|---|---|
| EMT | 3,133 | 4 | human | TGF β activated EMT in A539 cell line. | PMID: 32358524 |
| Mouse HSPC | 49,302 | 3 | mouse | Mouse_hematopoiesis with lineage information. | PMID: 31974159 |
| Veres | 18,099 | 8 | human | This dataset profiles in vitro differentiation of human pluripotent stem cells toward pancreatic endocrine lineages. | PMID: 31068696 |
| Ebdata | 18,204 | 5 | human | Human embryonic stem cells differentiated as embryoid bodies over a 27-day period. | PMID: 31796933 |
| HSPC | 9983 | 4 | huamn | The molecular and cellular dynamics of hematopoietic stem and progenitor cells (HSPCs) during inflammatory response. | https://doi.org/10.1101/2023.03.09.531881 |

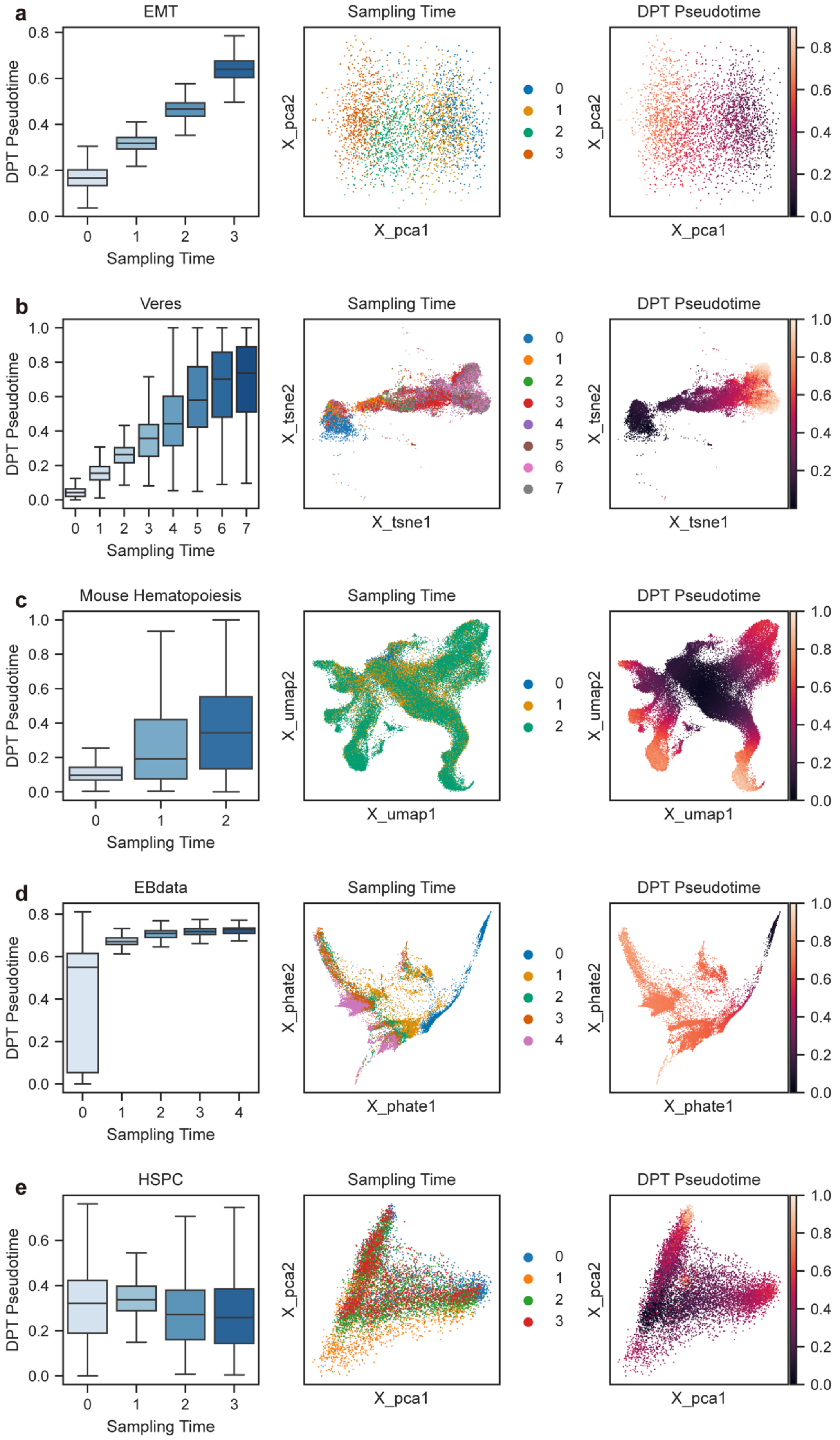
a
EMT
DPT Pseudotime
Sampling Time
Sampling Time
X_pca1
X_pca2
DPT Pseudotime
X_pca1
X_pca2
b
Veres
DPT Pseudotime
Sampling Time
Sampling Time
X_tsne1
X_tsne2
DPT Pseudotime
X_tsne1
X_tsne2
c
Mouse Hematopoiesis
DPT Pseudotime
Sampling Time
Sampling Time
X_umap1
X_umap2
DPT Pseudotime
X_umap1
X_umap2
d
EBdata
DPT Pseudotime
Sampling Time
Sampling Time
X_phate1
X_phate2
DPT Pseudotime
X_phate1
X_phate2
e
HSPC
DPT Pseudotime
Sampling Time
Sampling Time
X_pca1
X_pca2
DPT Pseudotime
X_pca1
X_pca2

**Supplementary Figure 1 | Assessment of pseudotime annotation quality across datasets.**
For each dataset, three complementary visualizations are shown. **Left**: distribution of DPT pseudotime values across experimental sampling time points, visualized as boxplots. High-quality pseudotime annotations (**a-c**) are expected to exhibit an approximately uniform spread within each sampling time and a monotonic increase in median pseudotime with later sampling times. **Middle**: two-dimensional visualization of the cell embeddings, colored by experimental sampling time. **Right**: the same embedding visualization, colored by pseudotime.

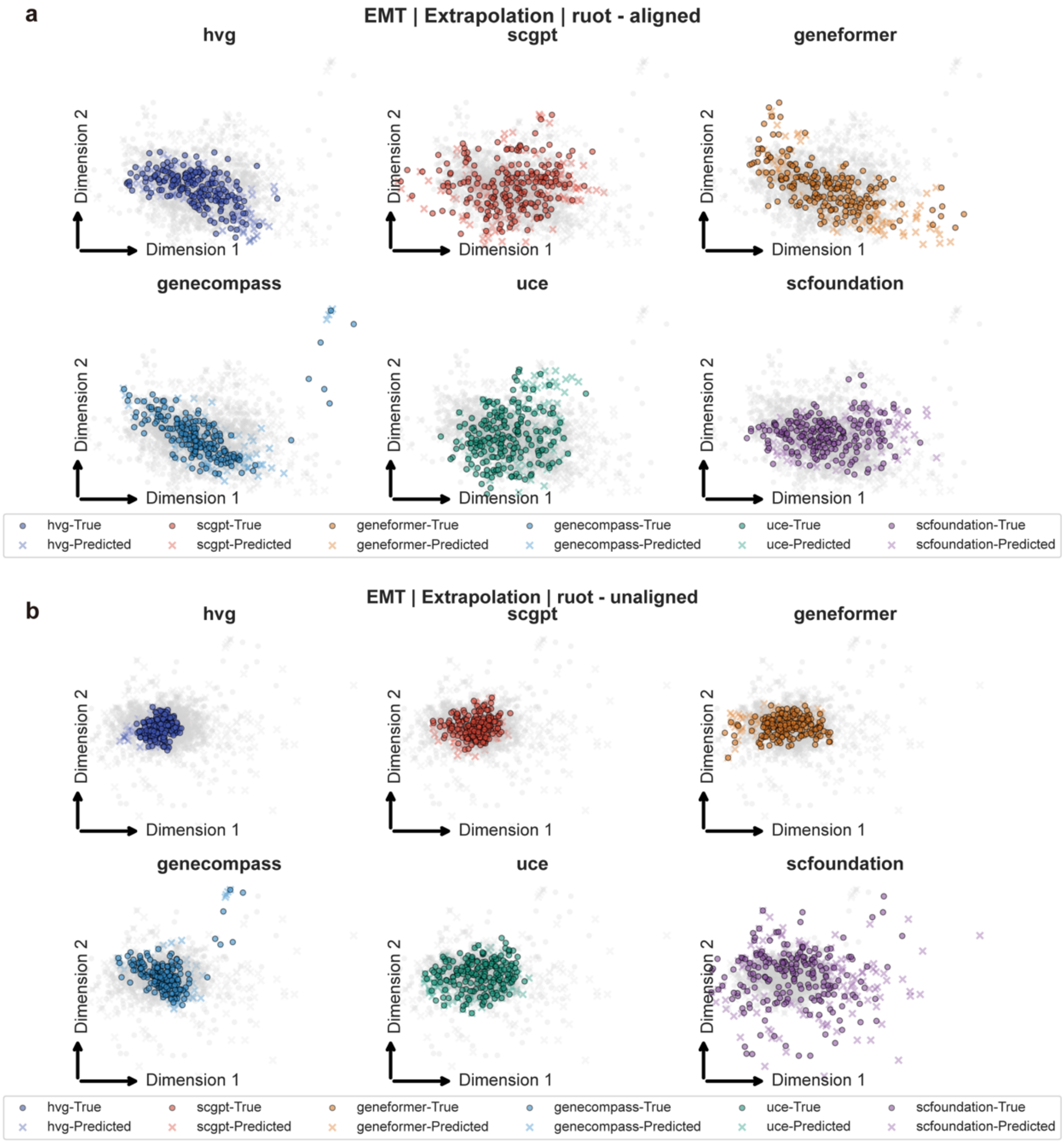


**Supplementary Figure 2 | Comparison of aligned versus unaligned embeddings for EMT dataset (extrapolation task).**

Scatter plot visualizations of observed and predicted cell states for six embedding methods (HVG, scGPT, Geneformer, GeneCompass, UCE, and scFoundation) on the EMT dataset in the extrapolation setting. Panels (a) and (b) show results with aligned and unaligned embeddings, respectively. Alignment places all embeddings in a shared latent space, reducing differences in scale and orientation and enabling fair comparison of predicted trajectories. Observed and predicted cell states are distinguished by marker shape.

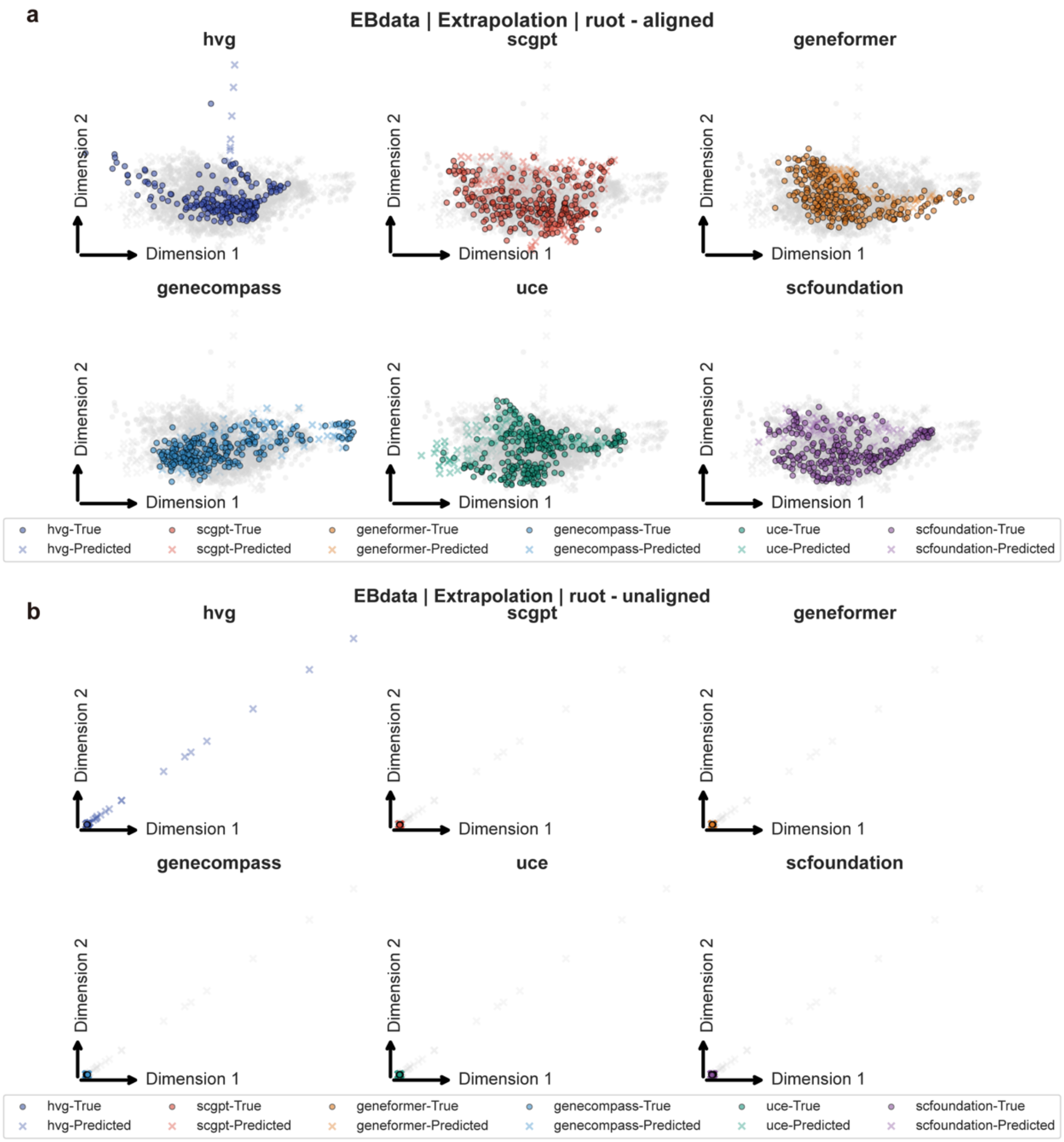


**Supplementary Figure 3 | Comparison of aligned versus unaligned embeddings for EBdata dataset (extrapolation task).**

Same as **Supplementary Figure 2**, for EBdata dataset.

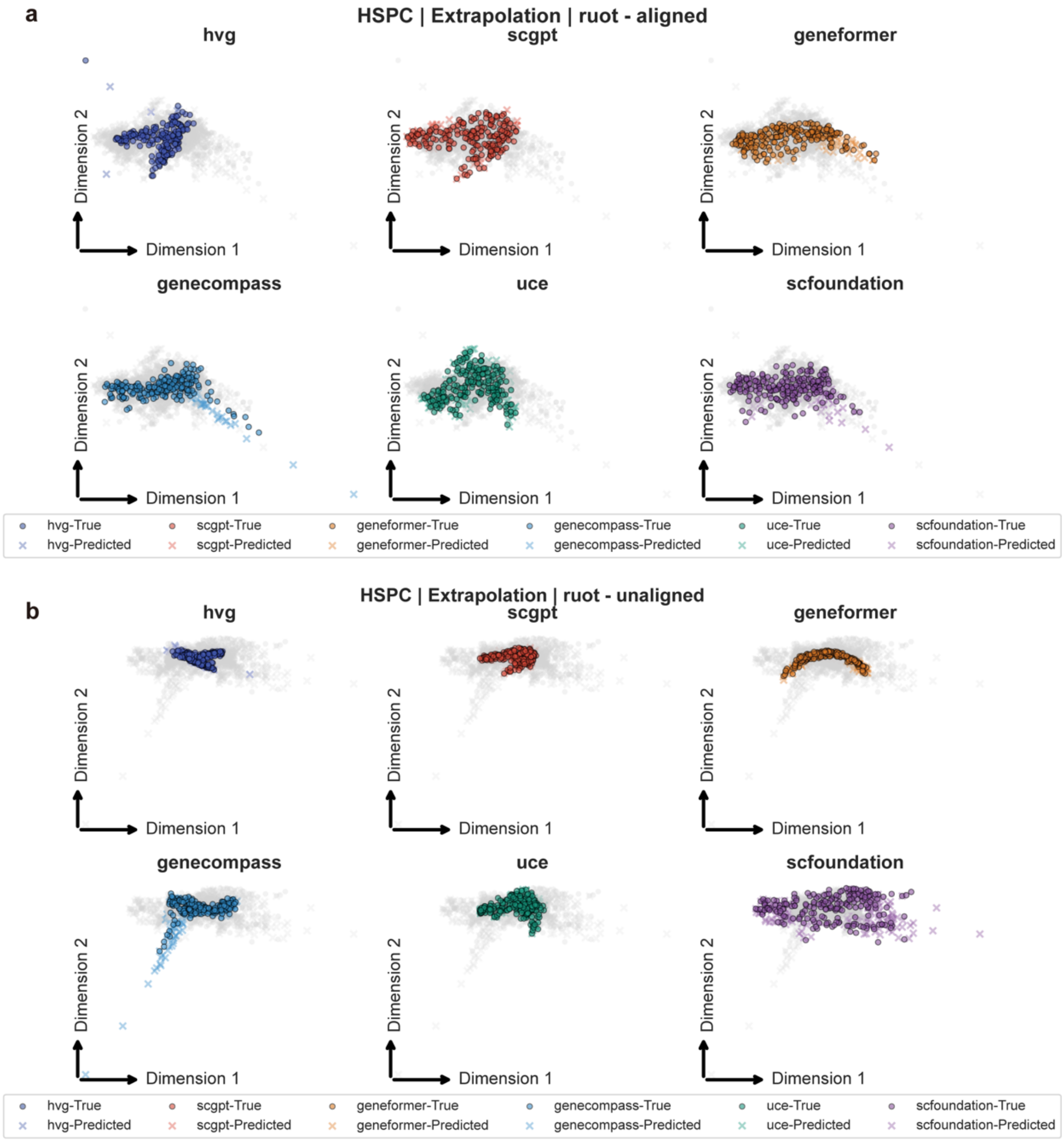


**Supplementary Figure 4 | Comparison of aligned versus unaligned embeddings for HSPC dataset (extrapolation task).**
Same as **Supplementary Figure 2**, for HSPC dataset.

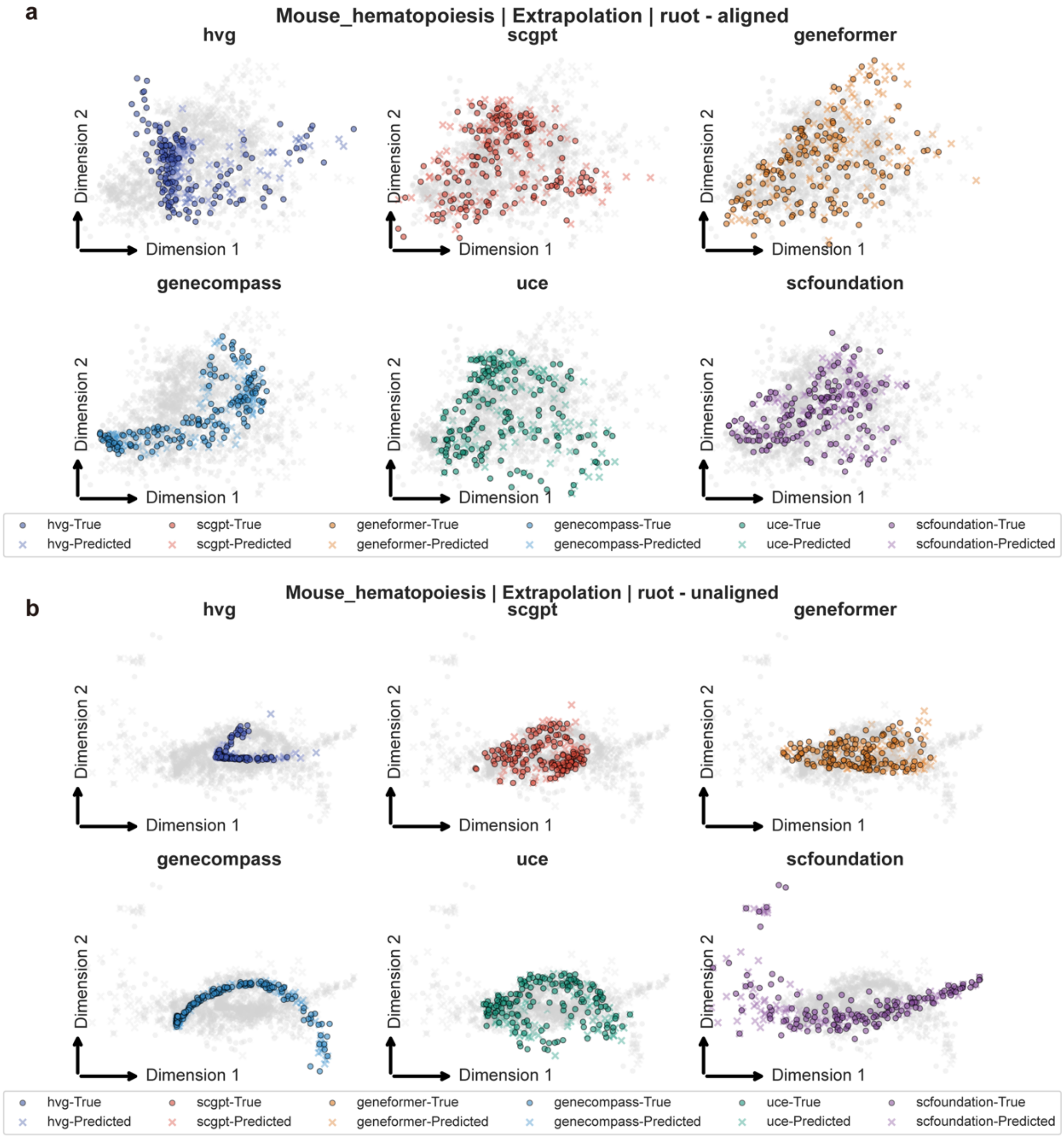


**Supplementary Figure 5 | Comparison of aligned versus unaligned embeddings for Mouse HSPC dataset (extrapolation task).**

Same as **Supplementary Figure 2**, for Mouse HSPC dataset.

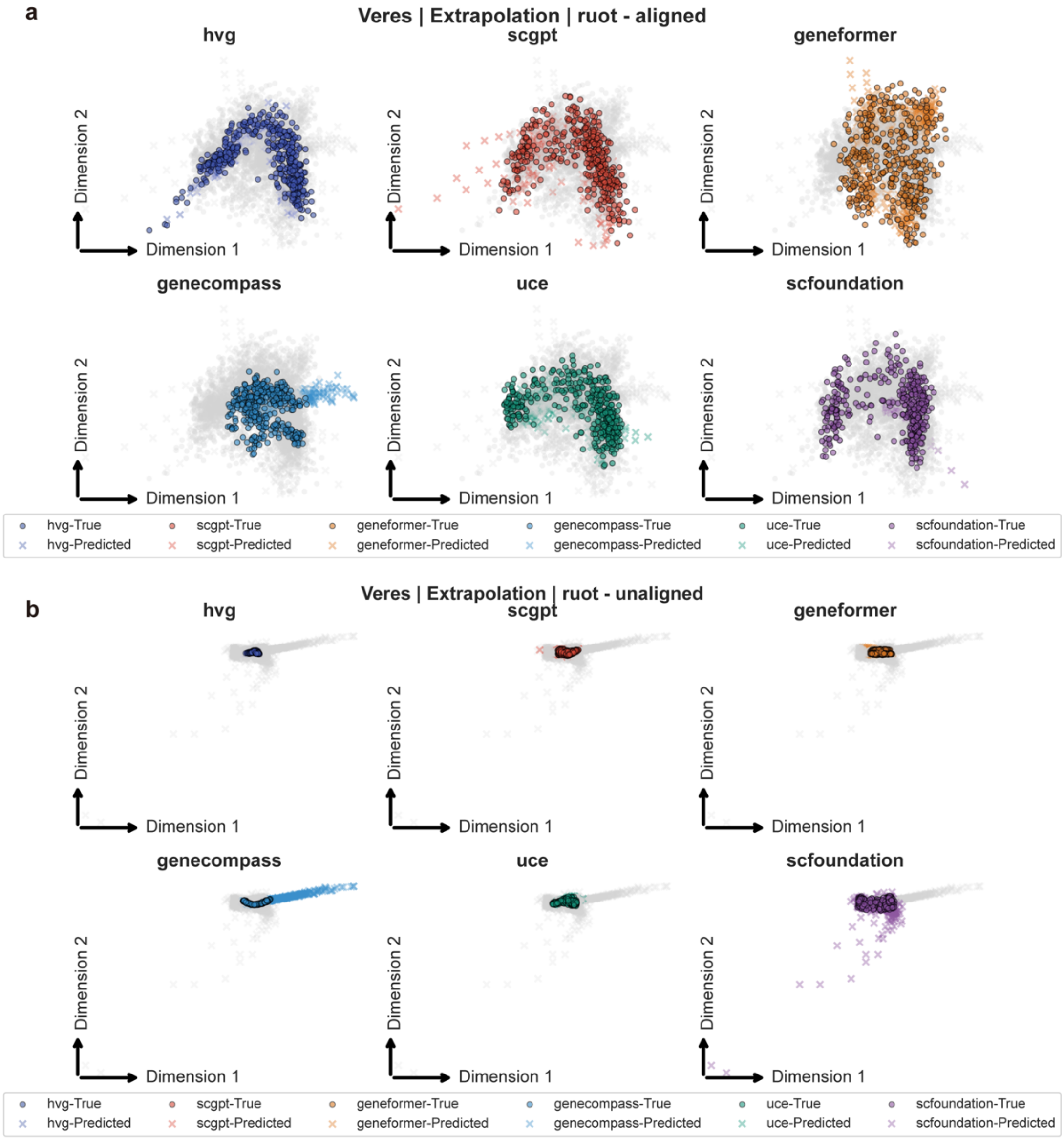


**Supplementary Figure 6 | Comparison of aligned versus unaligned embeddings for Veres dataset (extrapolation task).**
Same as **Supplementary Figure 2** for Veres dataset.

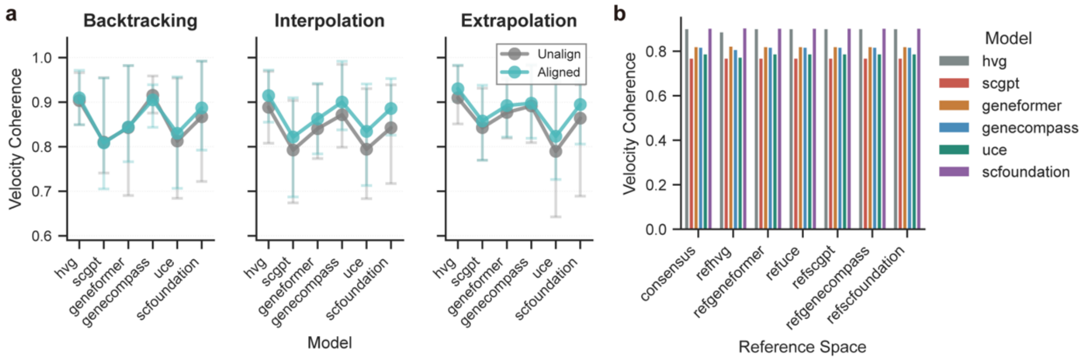


**Supplementary Figure 7 | Sensitivity of velocity coherence to alignment and reference space.**
**a,** Velocity coherence under aligned and unaligned evaluation settings.
**b**, Effect of reference space selection on velocity coherence, assessed by alternately treating each embedding as the alignment reference.
Across metrics, while absolute values vary modestly with alignment and reference-space choice, the relative ordering of embedders remains largely stable, consistent with the robustness observed for Wasserstein-1 distance in **Fig. 3**.

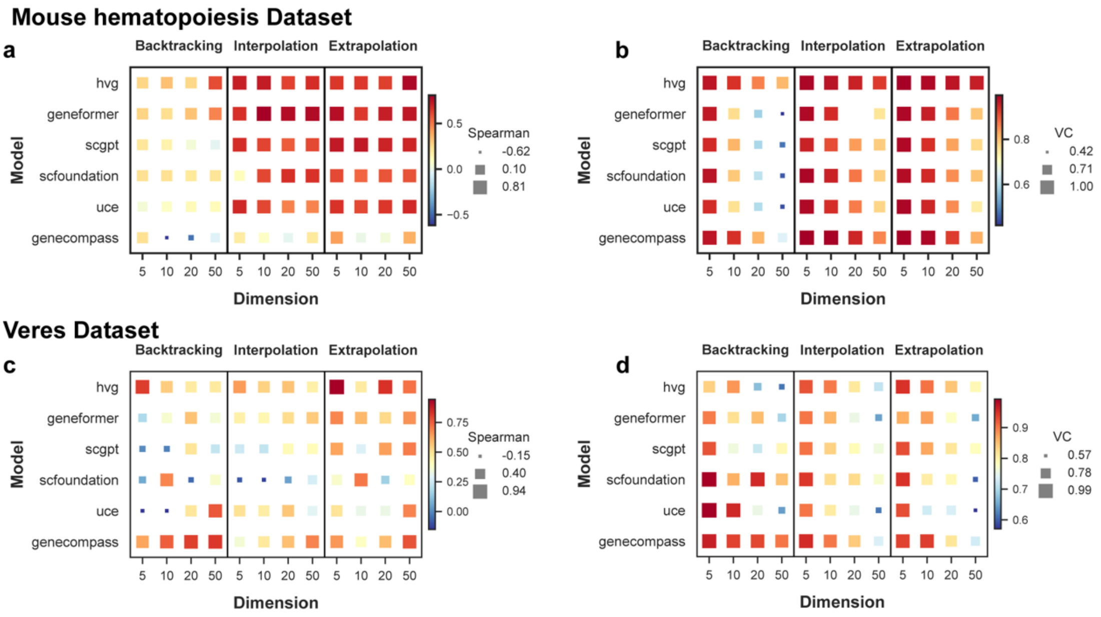


**Supplementary Figure 8 | Sensitivity of pseudotime correlation and velocity coherence to latent dimensionality across datasets.**

Sensitivity analyses corresponding to **Fig. 3 d, e**, extended to other benchmark datasets.

**a-b,** Effect of latent dimensionality on pseudotime correlation for Mouse HSPC dataset across the three dynamical tasks. Heatmaps show the Spearman correlation between inferred pseudotime and the reference pseudotime as a function of the number of retained principal components (5, 10, 20, and 50).

**c–d**, Effect of latent dimensionality on local velocity coherence across datasets and tasks. Heatmaps show velocity coherence as a function of latent dimensionality, with higher values indicating smoother and more self-consistent local velocity fields.

Across datasets, relative performance trends across embedders are largely preserved as latent dimensionality varies, although the dimensionality yielding optimal performance can be embedding- and dataset-specific.

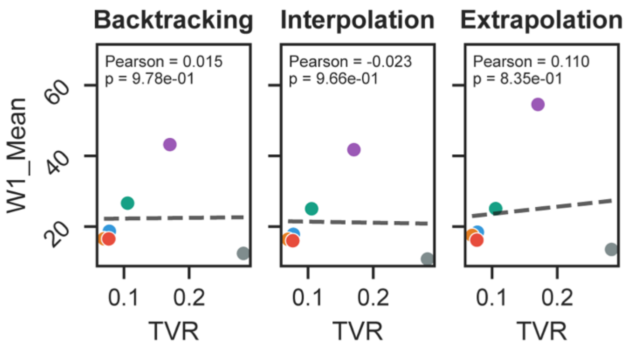


**Supplementary Figure 9 | Relationship between TVR and distributional reconstruction in the EMT dataset.**

Scatter plot showing the relationship between time variance ratio (TVR) and Wasserstein-1 (W1) distance in the EMT dataset across embeddings and dynamical task settings. Each point corresponds to one embedding–task combination, with TVR quantifying the fraction of time-associated variation preserved in the embedding space and W1 measuring the discrepancy between predicted and observed cell-state distributions. In contrast to the negative association observed in most datasets (Fig. 4b), the EMT dataset exhibits a weak or inconsistent relationship between TVR and W1. In particular, several foundation model embeddings display strong temporal compression (low TVR), with time points nearly co-localized in the embedding space, yet achieve low W1 distances. In this regime, distributional prediction becomes comparatively trivial because observed and target states are already close, illustrating a degenerate setting in which low W1 does not necessarily reflect faithful recovery of temporal dynamics

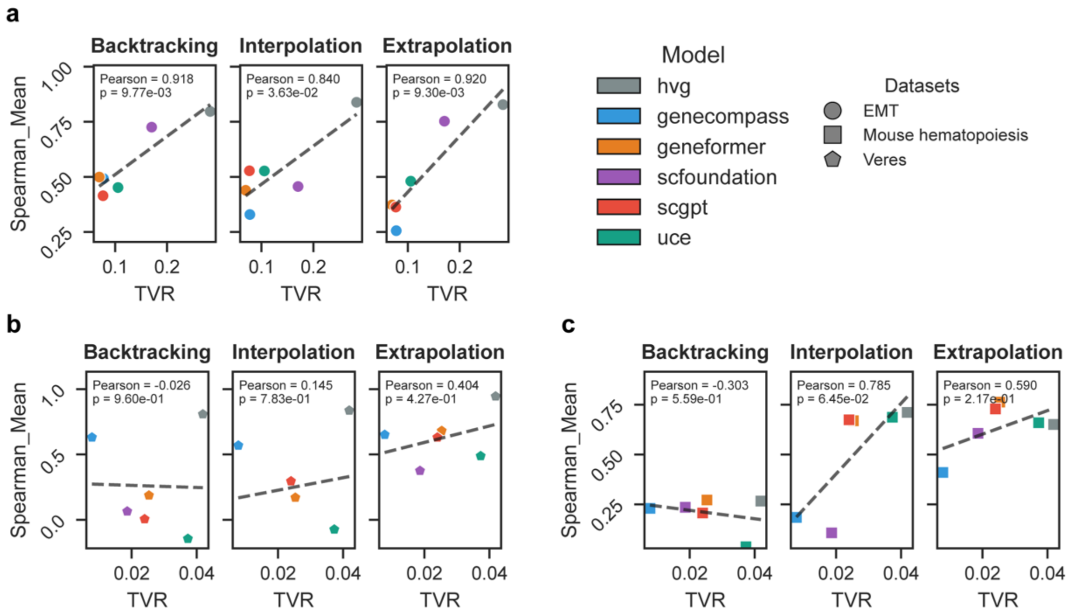


**Supplementary Figure 10 | Relationship between TVR and pseudotime correlation.** Scatter plots showing the relationship between time variance ratio (TVR) and pseudotime correlation across multiple datasets, embeddings, and task settings. Each point corresponds to one embedding–dataset–task combination. Pseudotime correlation is quantified as the Spearman rank correlation between inferred pseudotime and the reference pseudotime (Methods). TVR measures the fraction of time-associated variation preserved in the embedding space. The relationship between TVR and pseudotime correlation is heterogeneous across datasets and tasks. In several settings, embeddings with higher TVR exhibit improved pseudotime correlation, consistent with better preservation of global temporal ordering, whereas embeddings with very low TVR frequently show degraded ordering due to compression of time-structured variation.

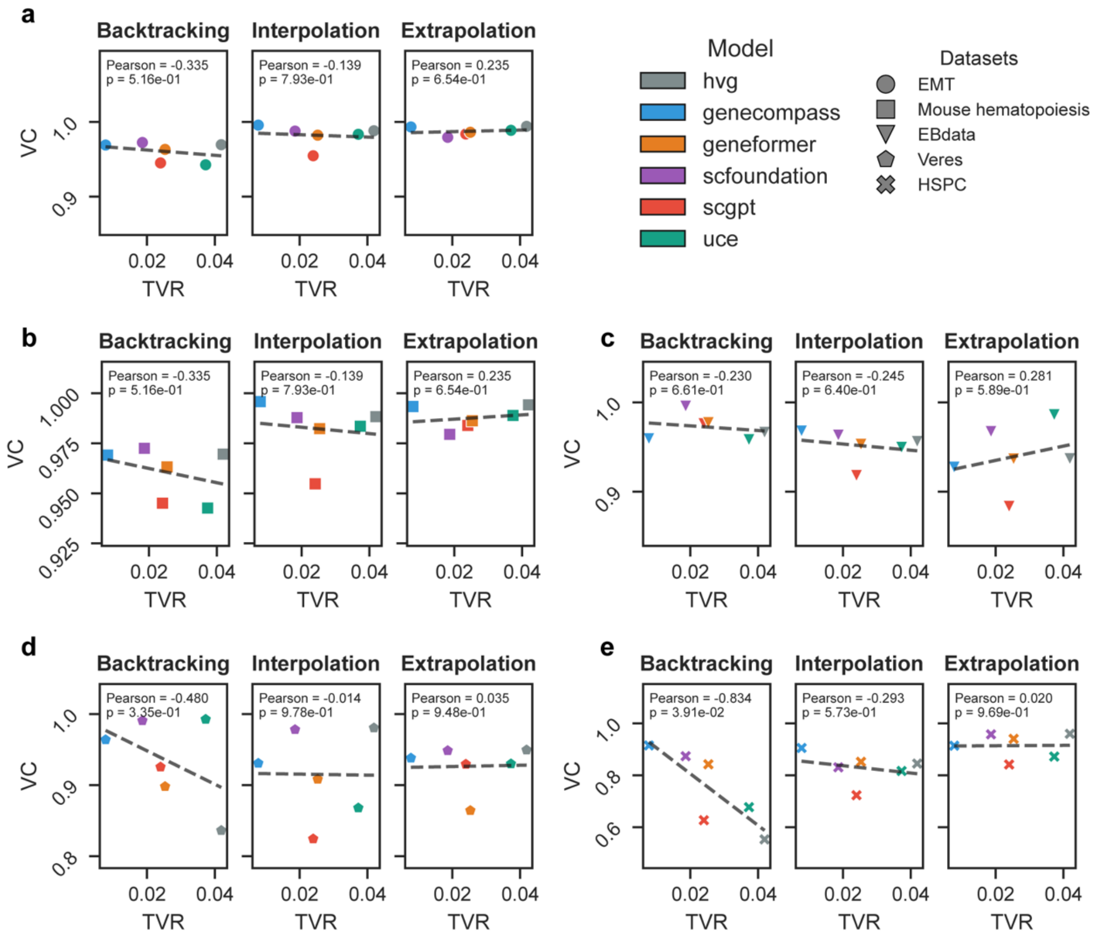


**Supplementary Figure 11 | Relationship between TVR and Velocity Coherence.**
Scatter plots showing the relationship between time variance ratio (TVR) and local velocity coherence across multiple datasets, embeddings, and task settings. Each point represents one embedding–dataset–task combination. Velocity coherence quantifies the agreement of inferred velocity vectors within local cell neighborhoods (Methods). In several settings, embeddings with higher TVR exhibit improved local velocity coherence, suggesting that excessive compression of time-associated variation can impair recovery of coherent local dynamics in some settings.

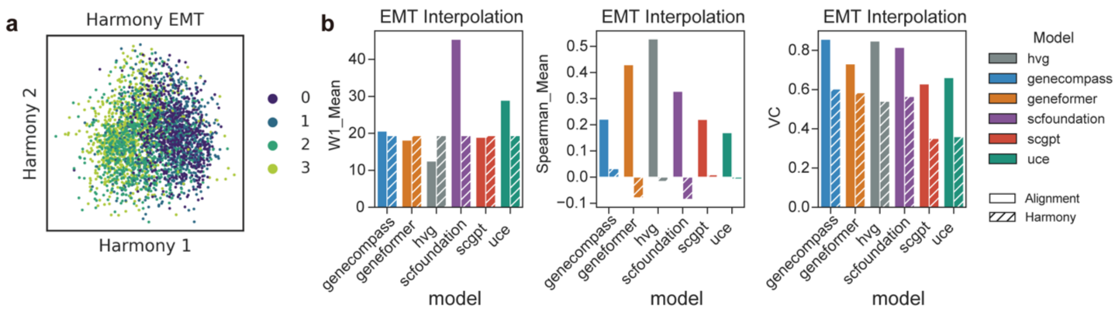


**Supplementary Figure 12 | Impact of explicit batch correction on dynamical metrics in the EMT dataset.**

**a.** Illustration of Harmony-based batch correction applied to HVG–PCA embeddings.

**b.** Effects of batch correction on dynamical reconstruction metrics. Grouped bar plots show three metrics evaluated in the aligned latent space: Wasserstein-1 distance (distributional recovery; lower is better), pseudotime correlation (Spearman ρ; higher is better), and local velocity coherence (higher is better). For each metric, solid-colored bars indicate original embeddings, while hatched bars indicate embeddings after Harmony-based batch correction. Batch correction reduces time-structured variance, equalizing W1 across embeddings, but concurrently decreases pseudotime correlation and velocity coherence, illustrating the trade-off between removing temporal variation and preserving dynamical information.

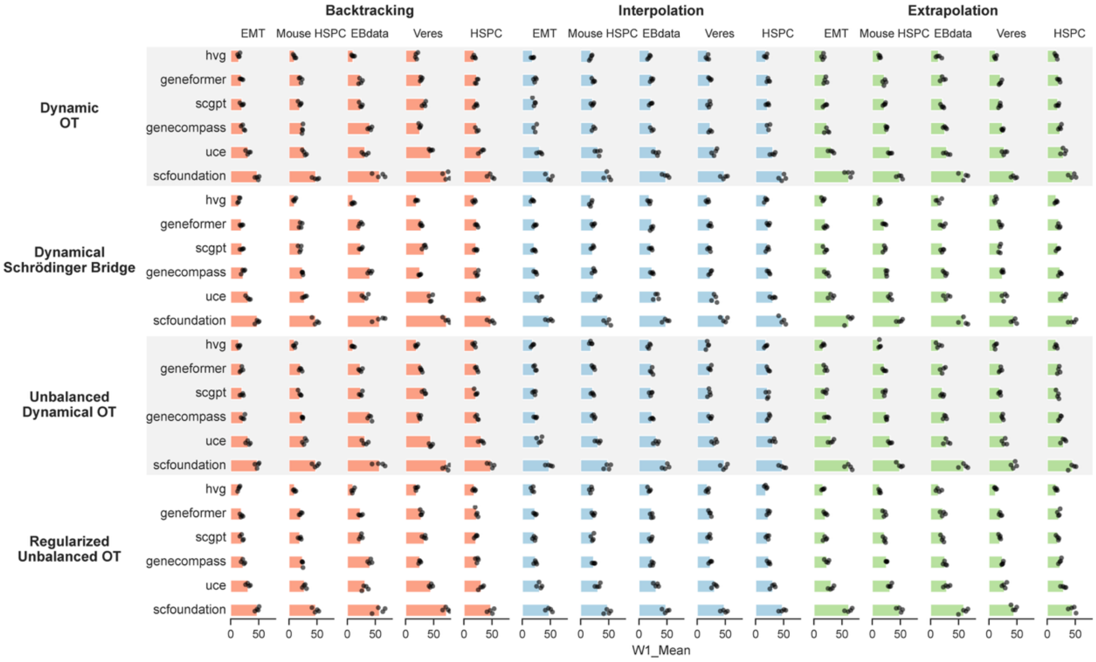


**Supplementary Figure 13 | Raw Wasserstein-1 distance across datasets, trajectory inference methods, and latent dimensionalities.**

Bar plots summarizing the Wasserstein-1 (W1) distance between predicted and observed cell-state distributions across all datasets and trajectory inference methods and task settings. Lower values indicate better distributional reconstruction. Each bar represents the mean W1 distance aggregated over latent dimensionality settings, while individual points denote results obtained under different latent space dimensions used for optimal transport–based dynamics inference (e.g., 5, 10, 20, and 50 dimensions). These raw metric values underlie the rank-based summaries shown in Fig. 5 and illustrate both absolute performance levels and variability across dimensionality choices.

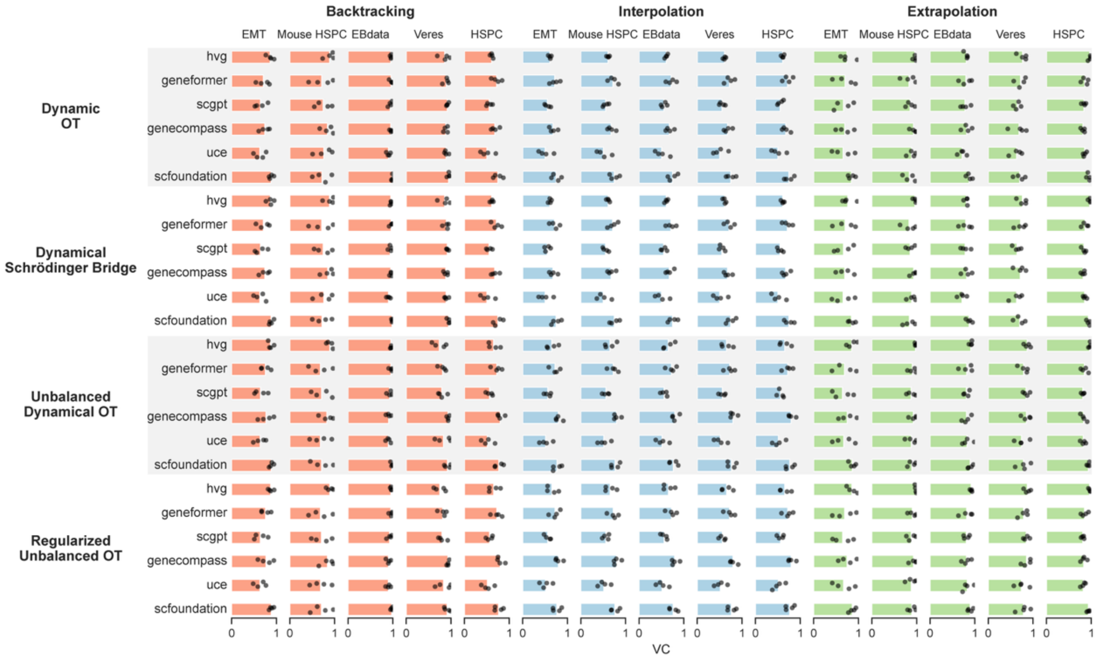


**Supplementary Figure 14 | Raw velocity coherence across datasets, trajectory inference methods, and latent dimensionalities.**

Bar plots summarizing local velocity coherence across all datasets, trajectory inference methods, and task settings. Higher values indicate stronger local directional consistency of inferred velocity fields. Each bar represents the mean velocity coherence aggregated over latent dimensionality settings, while individual points denote results obtained under different latent space dimensions used for optimal transport–based dynamics inference (e.g., 5, 10, 20, and 50 dimensions). These raw metric values underlie the rank-based summaries shown in Fig. 5 and illustrate both absolute performance levels and variability across dimensionality choices.
